# Oscillatory Control of Cortical Space as a Computational Dimension

**DOI:** 10.1101/2025.06.24.661361

**Authors:** Zhen Chen, Scott L. Brincat, Mikael Lundqvist, Roman F. Loonis, Melissa R. Warden, Earl K. Miller

## Abstract

Flexible cognition depends on the ability to represent and apply relevant information to the current task at hand. This allows the brain to interpret sensory input and guide behavior in a context-dependent manner. Recent work has proposed *Spatial Computing* as a mechanism for this flexibility, suggesting that task-related signals organize information processing through spatial patterns of oscillatory activity across the cortical surface. These patterns are proposed to act as “inhibitory stencils” that constrain where sensory-related information (the “content” of cognition) can be expressed in spiking activity. Here, we provide a comprehensive empirical test of Spatial Computing using multi-electrode recordings from the lateral prefrontal cortex in non-human primates performing a range of cognitive tasks (object working memory, sequence working memory, categorization). We found that alpha/beta oscillations encoded task-related information, were organized into spatial patterns that changed with task conditions, and inversely correlated with the spatial expression of sensory-related spiking activity. Furthermore, we found that alpha/beta oscillations reflected misattributions of task conditions and correlated with subjects’ trial-by-trial decisions. These findings validate core predictions of Spatial Computing suggesting that oscillatory dynamics not only gate information in time but also shape where in the cortex cognitive content is represented. This framework offers a unifying principle for understanding how the brain flexibly coordinates cognition through structured population dynamics.

## Introduction

A hallmark of cognition is its flexibility, the ability to adjust thoughts and actions according to current goals, rules, and contexts. This flexibility depends on more than just storing information. It requires the brain to control the expression of relevant information, when it should be accessed, and how it should be used. Central to this capacity is the use of learned information to shape how sensory inputs are interpreted and how behavior is guided.

Recent work has proposed a novel principle—*Spatial Computing*—as a potential mechanism for flexible context-dependent control ^1,2^. The core idea is that the brain achieves flexible control by using oscillatory dynamics to control neural activity across the cortical surface. In essence, cortical space is utilized as a computational dimension. In this framework, internally-generated signals reflecting goals and context are carried by oscillatory activity, particularly alpha/beta-band oscillations (10–30 Hz). By exerting inhibition, these low-dimensional mesoscale patterns modulate local neuronal excitability and determine which high-dimensional populations of neurons on the microscale can be activated in a given context. This shapes where in the cortex sensory-related spiking activity can occur. Rather than relying solely on fixed anatomical pathways, the brain dynamically organizes information flow through spatially structured oscillatory patterns. In this way, Spatial Computing offers a mechanism for how sensory information is processed and organized in a context-dependent manner.

This framework provides a mechanistic explanation for flexible control of cognition. For example, it helps explain how low-dimensional subspaces encoding different task variables can emerge, allowing the brain to reduce interference and flexibly switch between representations ^3,4^. It also accounts for how nonlinear mixed selectivity of neurons arises from the interaction of cognitive control and stimulus features ^5–8^. By dynamically modulating local neuronal excitability, alpha/beta oscillatory patterns allow the same neural populations to support different cognitive operations depending on context.

Despite these advances, several critical questions about Spatial Computing remain unanswered. First, the theory proposes that different neural signals serve distinct roles. Oscillatory patterns are proposed to reflect information learned to complete any given goal-directed task, which we refer to as “task information”. In contrast, spiking activity primarily represents bottom-up sensory inputs, the “*contents”* of cognition. However, little work has been done to directly compare the information carried by each signal type at the *population* level. Second, although Spatial Computing proposes that task information is reflected in alpha/beta oscillations, it is still unknown whether the spatial patterns of alpha/beta power reorganize with differing task conditions. It is also unknown whether any such reorganization between task conditions scales with cognitive demands, i.e. stronger cognitive demands result in larger task-based reorganization. Third, while the theory posits that alpha/beta regulates where sensory information is expressed, no direct test has shown that spiking-based sensory representation is spatially gated by alpha/beta patterns across the cortical space. Finally, although Spatial Computing is hypothesized to support flexible cognition and behavior, there is little direct evidence that alpha/beta or gamma (30–150 Hz) oscillations predict trial-by-trial decisions or reflect behavioral errors. Together, these open questions point to the need for a more comprehensive empirical test of Spatial Computing Theory.

We analyzed multi-electrode recordings from the lateral prefrontal cortex as non-human primates (NHPs) performed various cognitive tasks requiring working memory (WM), sequencing, and/or categorization. We formalized five core predictions of Spatial Computing and tested them across different cognitive demands. Our results show that alpha/beta patterns encode task information, flexibly reorganize with task conditions, spatially govern where sensory information is expressed in spiking activity, and closely track behavioral performance. These results provide empirical support for Spatial Computing Theory. They highlight its potential as a unifying framework for understanding how the brain flexibly coordinates cognition through structured population dynamics.

## Results

### Predictions of Spatial Computing Theory and Tasks

We tested five basic predictions of *Spatial Computing Theory*: 1. Alpha/beta oscillations primarily convey information reflecting cognitive control and task. 2. Spiking activity conveys both sensory (e.g., stimulus identity) and task information. 3. Alpha/beta power is spatially organized by task condition across the surface of cortex. 4. Sensory information is spatially anti-correlated with shifting alpha/beta patterns reflecting task information. 5. Alpha beta oscillations reflect N s’ behavioral performance and trial-by-trial choices. As downstream effectors of cognitive control and structured by alpha/beta, spiking and associated gamma are expected to correlate with behavior as well. **Figure 1A** illustrates this framework, where alpha/beta patterns carry task information and are dynamically shaped by the current task condition. These alpha/beta patterns act as “inhibitory stencils” which constrain where higher levels of spiking occur. This allows the stimulus-driven sensory information to be expressed only in regions where alpha/beta is weak. In essence, cortical space becomes a computational dimension. Alpha/beta patterns sculpt the spatial landscape of neural activity, thereby supporting context-dependent information processing.

**Figure 1.**
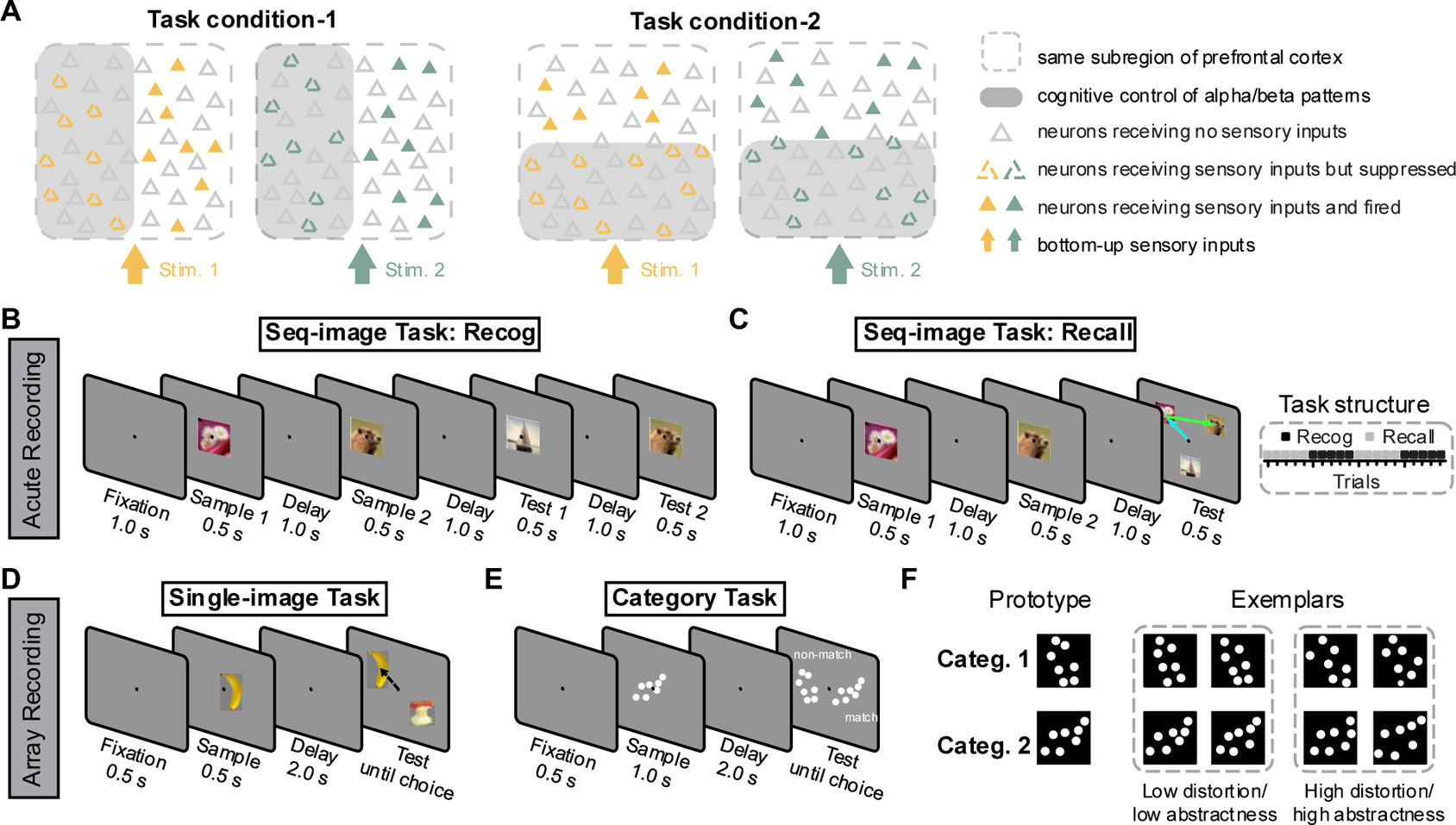
Schematic overview of Spatial Computing and task designs. (A). Conceptual framework of Spatial Computing Theory. Alpha/beta oscillations convey cognitive control signals reflecting learned task information, and their spatial pattern gates where sensory information (“content”) can be expressed by spiking. Shaded grey areas indicate stronger alpha/beta power where sensory information is suppressed while unshaded areas indicate lower alpha/beta power where sensory information is expressed by spiking. (B). “Recognition” rule version of Seq-image Task showing a sequence of sample images followed by a delay. During the test phase, NHPs judged whether a probe sequence matched the sample sequence. (C). “Recall” rule version of Seq-image Task where, during the test phase, NHPs made saccades to the remembered objects in their original order. The two rule versions alternated in trial blocks (right). (D). Single-image task where a single image was shown and, after a delay, NHPs made a saccade to the remembered object. (E). Category-match task where NHPs categorized exemplar dot patterns into two learned categories. (F). The exemplar dot patterns were generated by distorting two prototype patterns (left). Distortion levels varied the similarity between exemplars and prototypes, controlling the degree of abstractness (low vs. high).

To test these predictions, we analyzed multi-electrode neurophysiological recordings (local field potentials (LFPs) and single/multi-unit spiking activity) from the lateral prefrontal cortex (PFC) of NHPs (rhesus macaques) performing three cognitive tasks (**Figure S1**). In one of the tasks, the Seq-image Task ^9^, the NHPs performed a sequential working memory (WM) task involving two sample items. Two versions of the task were used. Both required working memory for two sample items and their order of appearance. They only differed in the rule that determined how NHPs reported these memories. In the ‘Recognition’ version, the NHPs indicated whether a subsequent sequence of test items was a match to the sample sequence (**Figure 1B**). In the ‘Recall’ version, NHPs were offered three test items and asked to saccade to the two seen earlier in the correct order (**Figure 1C**). The NHPs alternated between the two versions across blocks of trials. Thus, NHPs needed to maintain two types of task information: the order in which the two samples were presented on each trial, and the current rule that needed to be applied to the stored samples and test items.

The two other tasks involved working memory and categorization of individual images. The Single-image task ^10,11^ was a standard delayed match-to-sample working memory task (**Figure 1D**). In the Category Task ^12,13^, NHPs had to categorize dot-pattern stimuli into two learned categories, each defined by random distortions around a prototype (**Figure 1E1**). The level of distortion (“abstractness”) was varied by manipulating the similarity between the exemplars and their corresponding prototype (**Figure 1E2**). Thus, in this task, the individual dot exemplars were the sensory inputs to cognitive processing. Their inferred category was the learned task condition that determined which behavioral response was appropriate for each exemplar. Additionally, manipulating the level of exemplar distortion across trials allowed us to test different degrees of cognitive demands.

These three tasks allowed us to probe how sensory information (e.g., identity of sample items and exemplar dot patterns) and task information (e.g., rule, order, and category) are encoded by spiking and LFP dynamics. By examining the spatial distribution of these signals, we could directly test whether alpha/beta patterns are spatially organized, consistent with a “stencil” that organizes sensory information expressed by spiking. A complete description of the experimental design for each task can be found in the **Methods**. Trial-averaged power spectrograms for each dataset are provided in **Figure S2**, with additional figures and analyses available in the original publications that first reported these data ^1,9–14^. The spectrograms showed clear spectral peaks within the 10–30 Hz band, indicating our results reflect true alpha/beta oscillations, rather than broadband aperiodic effects ^15^.

### Predictions 1 and 2: Alpha/beta Reflects Task Information, Spiking Reflects Both Sensory and Task Information

To test these predictions, we quantified the amount of information reflecting the sensory stimuli and task conditions carried by each signal type. Rather than analyzing single electrodes or individual units, we developed a method to compute the percentage of explained variance (PEV) by each factor across the entire population of units or LFPs (see **Methods**). For each factor, we computed an axis in population activity (spike rates or LFP band power) space that optimally separated contrasted conditions. We then projected population activity onto this axis and quantified its PEV. This approach allowed us to capture the information encoded by population-level LFP power and spiking.

We began with the sensory information in the Seq-image Task—the identity of the two sample items to be remembered (**Figure 1B**). In line with previous studies ^16^, we found that spiking activity carried the most sensory information. Information increased sharply during each sample presentation (**Figure 2A**). There was some sensory information in gamma, but it was much lower than in spiking activity (**Figure S3**). Sensory information was minimal in alpha/beta power, in line with Prediction 1.

**Figure 2.**
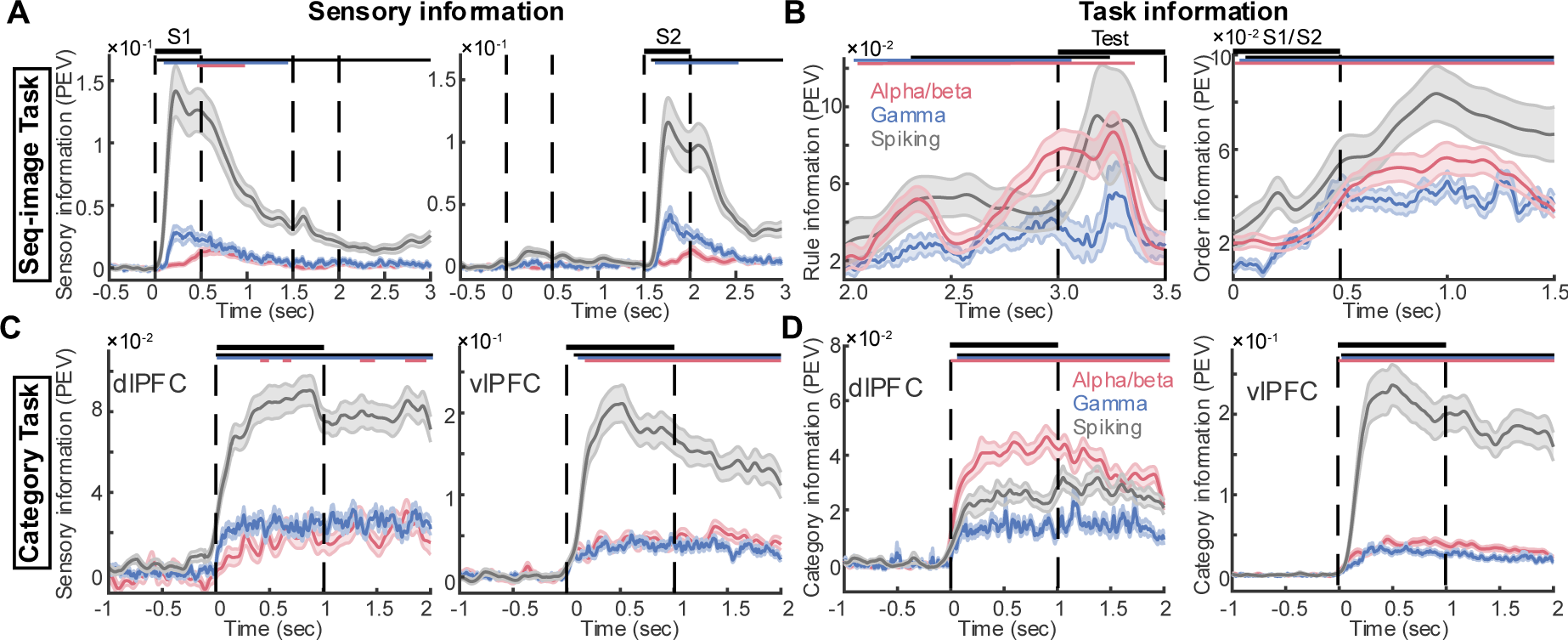
Alpha/beta power primarily carried task information, while spiking activity conveyed mixed sensory and task information. (A). Sensory information (quantified by PEV) conveyed by each signal type in the Seq-image Task, for sample 1 (left) and sample 2 (right). Vertical dashed lines indicated the sample presentation periods (S1 and S2). Horizontal thin lines on top reflect times of significant PEV values for the corresponding signal type using the same color code (p<0.001, one-sample t-test). Error bar: mean ± SE, n = 48 (2 rules × 24 sessions). (B). Task information conveyed by each signal type in the Seq-image Task. Left: PEV for rule (Recognition vs. Recall) during the second delay and test period. Right: PEV for sequence order (first vs. second order) during the sample presentation followed by delay. Error bar: mean ± SE, n = 24 sessions. Significance: p<0.001, one-sample t-test. (C). Sensory information for each signal type in the Category Task. Left: PEV for dlPFC. Right: PEV for vlPFC. Error bar: mean ± SE, n = 51 sessions. Significance: p<0.001, one-sample t-test. (D). Category information for each signal type in the Category Task. Left: PEV for dlPFC. Right: PEV for vlPFC. Error bar: mean ± SE, n = 51 sessions. Significance: p<0.001, one-sample t-test. See Fig. S2 for corresponding paired comparisons between signal types.

We then assessed task condition encoding by measuring task information conveyed about the *rule* being performed (i.e., Recognition vs. Recall) during the delay period preceding test onset. Throughout the delay, all signals carried task information reflecting the current rule but alpha/beta and spiking carried more than gamma (**Figure 2B**, left; see **Figure S3** for paired comparison). Note the ramping up of alpha/beta information reflecting the rule. It peaked near the end of the delay and plateaued through the presentation of the test items when the rule needed to be applied. In addition to the rule, we also examined the sample sequence order (i.e., first vs. second sample item), which was another type of task factor in this paradigm. We computed information reflecting the sequence presentation order (i.e., sample/delay 1 vs. sample/delay2), while holding constant the sample image shown (see **Methods**). Similar to information reflecting the rule, all signals carried sequence order information, with alpha/beta and spiking carrying more than gamma (**Figure 2B**, right; see **Figure S3** for paired comparison). These results showed that, in the Seq-image Task, task information was reflected mainly in alpha/beta power and spiking, while WM content was reflected primarily in spiking activity, consistent with Predictions 1 and 2.

We then turned to the Category Task. The sensory stimuli in this task were the individual category exemplars, while their inferred category memberships were learned task conditions that determined the appropriate behavior for each exemplar. For this dataset, we have extensive sampling of both the dorsolateral (dlPFC) and ventrolateral (vlPFC) prefrontal subregions. In both subregions, spiking activity dominated sensory information, while alpha/beta and gamma in vlPFC also carried a modest amount (**Figure 2C**, see **Figure S3** for paired comparison). By contrast, for category information, we observed a clear distinction between the two regions, as we discuss next.

Previous work has shown that the dlPFC is more associated with top-down processing, while vlPFC is relatively more driven by bottom-up sensory stimulation ^13,17–19^. Consistent with this, alpha/beta oscillations in dlPFC were the primary carriers of category information (**Figure 2D**, left; see **Figure S3** for paired comparison). By contrast, in vlPFC, spiking activity dominated (**Figure 2D**, right; see **Figure S2** for paired comparison). Importantly, these differences between dlPFC and vlPFC were not due to differences in overall spiking rate or recording quality between these two regions. Although vlPFC showed a brief (∼100 ms) elevation in mean spiking rate relative to dlPFC (**Figure S4**), their spiking rates were nearly identical through the rest of the trial. Moreover, dlPFC and vlPFC carried comparable amounts of information reflecting the behavioral response (choice of left vs. right test exemplar; **Figure S4**). This indicated that the observed differences between dlPFC and vlPFC arose from their functional distinctions. In dlPFC alpha/beta mainly reflected category, while in vlPFC spiking conveyed both category and sensory inputs more prominently. This dorsolateral/ventrolateral distinction suggested a gradient in the neural implementation of top-down versus bottom-up processing, which we return to in the Discussion.

For a further test of category, we examined the level of abstractness of the categories. The Category Task manipulated the level of abstractness by different degrees of distortions of the prototypes. For low distortions, the exemplars from the same category were more similar in appearance to one another and to their prototypes. Thus, they could be categorized by their bottom-up appearance, and required less abstraction. By contrast, at high distortion levels, exemplars from different categories could appear similar to each other. They thus had to be categorized by a task rule learned by the NHPs ^13^.

This revealed category information in alpha/beta was stronger for the high-abstractness exemplars (especially in dlPFC). By contrast, category information in spiking was stronger for the low-abstractness exemplars (in vlPFC). In dlPFC, alpha/beta (**Figure 3A,B**, left) and gamma (**Figure 3A,B**, center) power encoded significantly more category information under high-abstractness than under low-abstractness. These effects were greater than the modulation of dlPFC spiking activity by abstractness, which was itself not significant (**Figure 3A,B**, right). In vlPFC, by contrast, spiking activity carried more category information under low-abstractness conditions than under high abstractness (**Figure 3C,D**, right), while alpha/beta and gamma showed the opposite direction of modulation by abstractness level (**Figure 3C,D**, left and center). Thus, when exemplars were heavily distorted (high-abstractness), dlPFC alpha/beta plays a dominant role in representing the relevant category through top-down mechanisms. Conversely, when exemplars were easily recognized (low-abstractness), vlPFC spiking could represent the category more directly through bottom-up mechanisms. This is consistent with Predictions 1 and 2.

**Figure 3.**
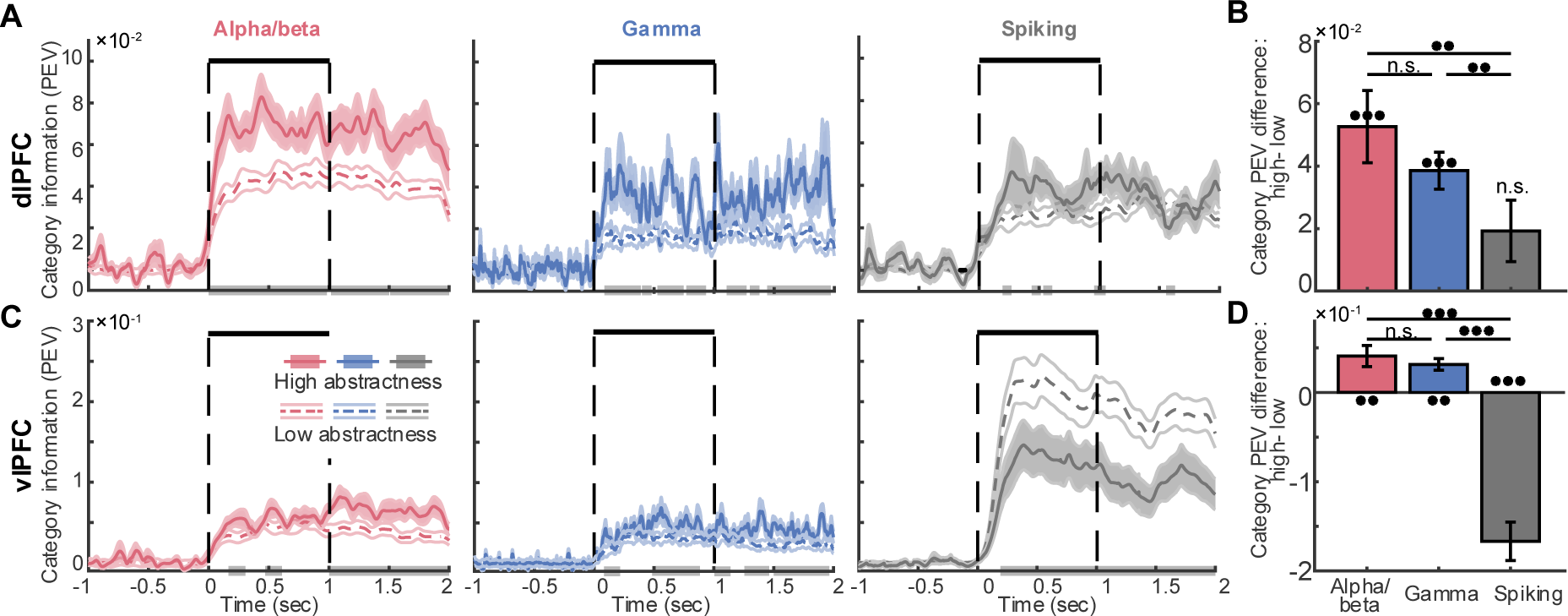
Category encoding varies with abstractness level. (A). Category information (PEV) by alpha/beta (left), gamma (middle), and spiking (right) in dlPFC, shown separately for high- (solid lines) and low-abstractness (dashed lines) conditions. Horizontal grey bars on the x-axis represented significant PEV differences between high- and low-abstractness (p<0.05, two-sample t-test). Error bar: mean ± SE, n = 51 sessions. (B). Cumulative difference in category PEV (high-minus low-abstractness) across time in dlPFC, shown for each signal type. *** p<0.001, **: p<0.01, n.s.: not significant, one-sample t-test for each signal type and Wilcoxon rank-sum test across signals. Error bar: mean ± SE, n = 51 sessions. (C). Same as (A) but for vlPFC. (D). Same as (B) but for vlPFC.

Overall, these findings support Predictions 1 and 2 of Spatial Computing. They showed that alpha/beta (especially in dlPFC) strongly reflected learned task information, while spiking reflected both sensory and task information. Task information carried by alpha/beta (in dlPFC) was greater when tasks required greater cognitive demands. The distinction between dlPFC and vlPFC aligned with their respective bias toward top-down vs. bottom-up processing.

### Prediction 3: Alpha/beta Power is Spatially Organized by Task Condition Across the Surface of Cortex

Spatial Computing theory further predicts that alpha/beta oscillations form spatial patterns across the cortical surface that reflect task information and reorganize with changing task conditions. A critical component of this prediction is that, for each distinct task condition, alpha/beta power should exhibit spatial clustering, rather than being spatially unstructured. Because the Category Task and Single-image Task used multi-electrode Utah arrays (see **Methods**), we could test this prediction directly with these datasets.

To quantify the spatial clustering of alpha/beta power within each array, we computed Moran’s I ^20–22^, a common measure of the overall strength of spatial autocorrelation (see **Methods**). We compared the observed values to those obtained from spatially shuffled versions of the same patterns. If alpha/beta power is spatially clustered, its spatial autocorrelation should be significantly positive. By contrast, the shuffled patterns, lacking spatial structure, should yield autocorrelation values near zero.

We found that in both dlPFC and vlPFC, Moran’s I values for the original alpha beta patterns were significantly greater than those of the shuffled patterns, which were around zero (**Figure 4A,B**, left). This effect held for both low and high levels of abstractness of the Category Task, as well as for the Single-Image Task (**Figure S5A,B**, left). We confirmed this spatial clustering with an alternative metric by computing the absolute power differences between neighboring electrodes (see **Methods**). This yielded consistent results: shuffled patterns exhibited significantly larger power differences between nearest neighbors compared to the actual observed alpha/beta patterns **(Figure S5A,B**, right; **S5C,D**). These results indicated that alpha/beta power was spatially organized across the cortical surface, as predicted by Spatial Computing Theory.

**Figure 4.**
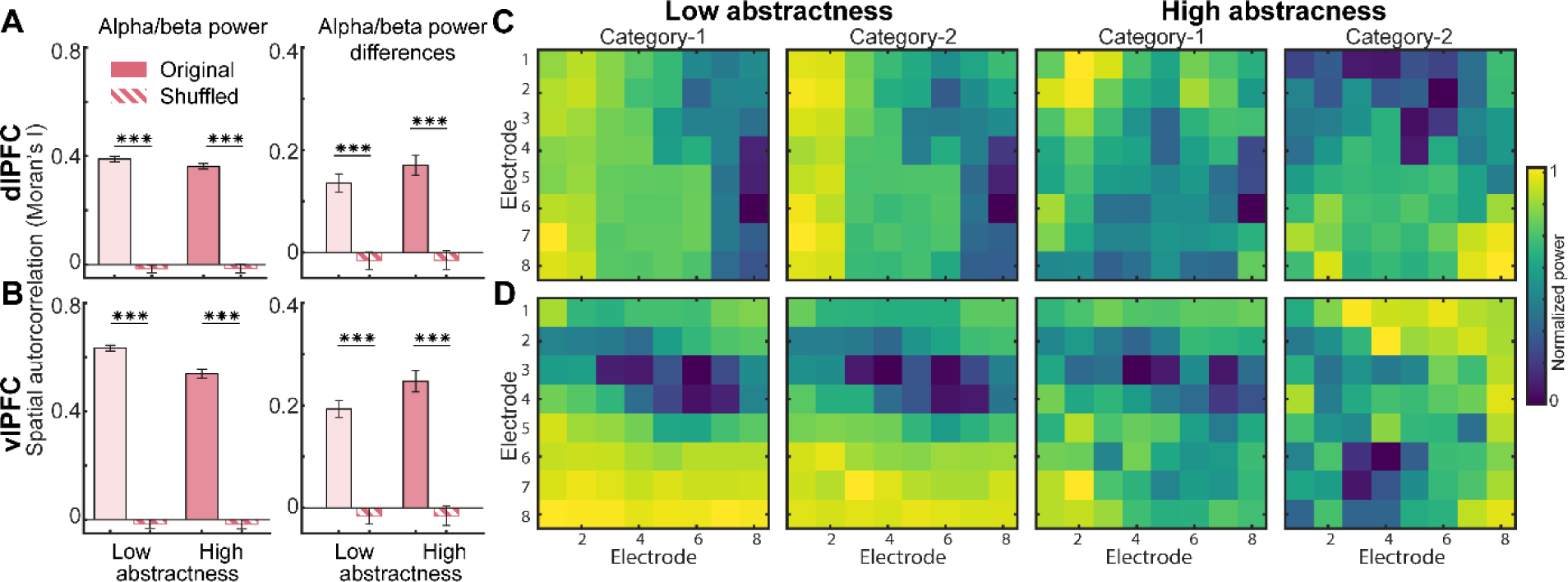
Alpha/beta power is spatially organized across the surface of cortex. (A). (Left). Spatial autocorrelation, measured by Moran’s I, of the trial-averaged patterns of alpha/beta power during sample presentation in dlPFC (solid bars), for low and high abstractness of the Category Task. As a control, Moran’s I for patterns randomly shuffled across space is shown (hatched bars). (Right). Same as left but for alpha/beta power differences between category-1 and category-2. *** p<0.001, two-sample t-test. Error bar: mean ± SE, n = 51 sessions. (B). Same as (A) but for vlPFC. (C). Example spatial patterns of alpha/beta power averaged across trials, shown for two categories under low- and high-abstractness in one session from dlPFC. (D). Same as (C) but for vlPFC.

Spatial Computing theory predicts spatial organization not only of overall alpha/beta *power*, but also of the task information it carries. That is, alpha/beta at nearby sites should “prefer” the same category. To quantify this, we computed the signed power difference between categories (Category-1 – Category-2) at each electrode. This is a measure of category selectivity that reflects both its category preference (sign) and strength (magnitude). We then quantified their spatial autocorrelation with Moran’s I and compared them against spatially shuffled controls. These category selectivity maps showed significantly positive spatial autocorrelation, whereas shuffled maps were near zero (**Figure 4A,B**, right). These results reflected contiguous clusters of electrodes that consistently favor one category over the other. Thus, alpha/beta power formed patterns that were spatially organized and category-selective. Moreover, further spatial correlation analysis confirmed that these patterns reconfigured with category (i.e., lower between-category correlation than within-category, **Figure S5E-H**). Example spatial maps in **Figure 4C** and **4D** illustrated this spatially structured organization of alpha/beta patterns with increasing category differentiation under high-abstractness in dlPFC and vlPFC respectively. These results are consistent with Prediction 3.

### Prediction 4: Sensory Information is Spatially Anti-correlated with Shifting Alpha/Beta Patterns Reflecting Task Information

Alpha/beta patterns are proposed to spatially regulate the expression of sensory information carried by spiking. This spatial regulation is expected to shift with changing task conditions. **Figure 5A** shows a schematic of this hypothesis: In each task condition, areas of high alpha/beta power exhibit lower sensory information carried by spiking, whereas areas with low alpha/beta power show higher spiking sensory information. Moreover, the spatial pattern of alpha/beta, and thus the location of spiking sensory expression, varies across task conditions.

**Figure 5.**
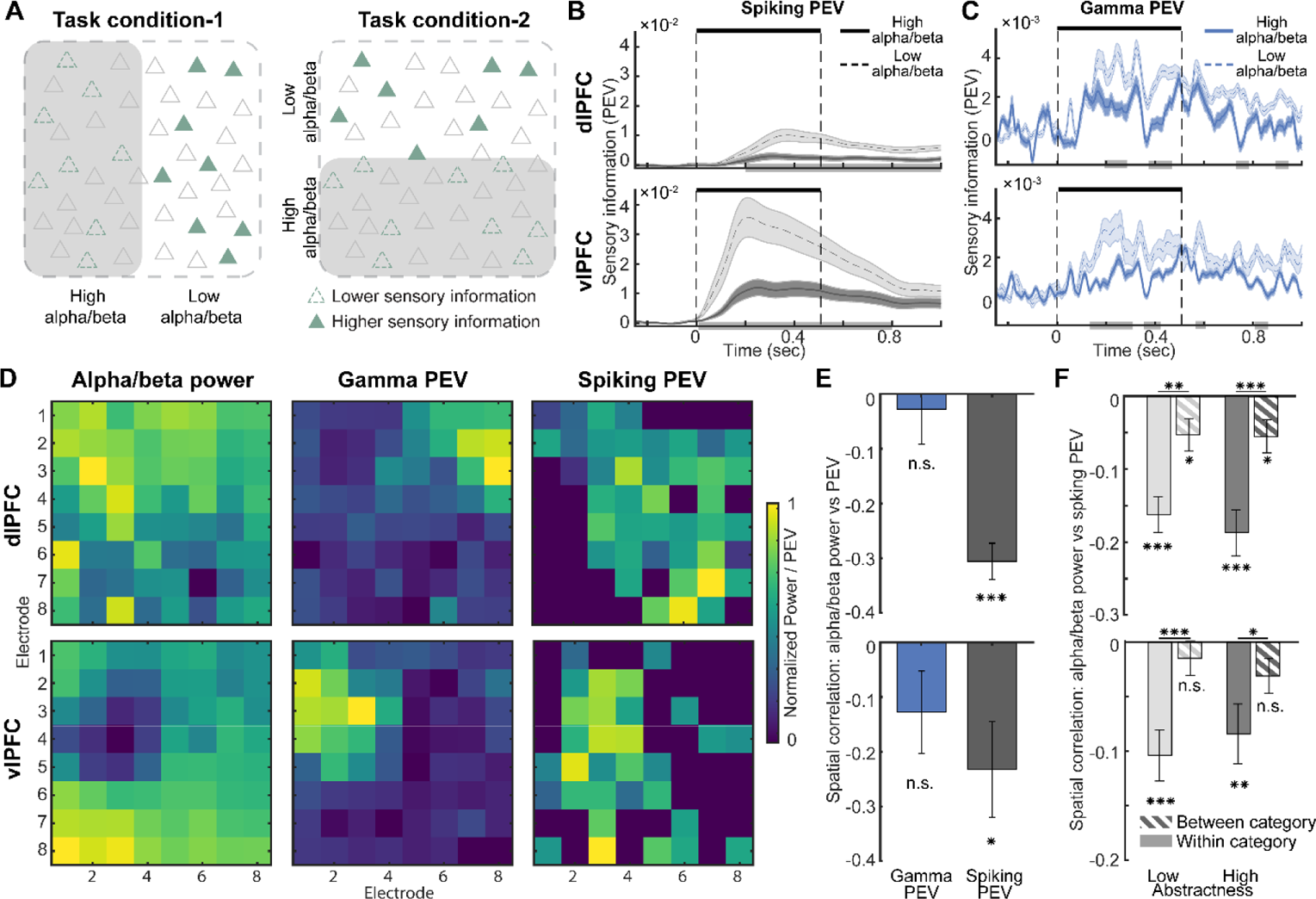
Alpha/beta patterns are spatially anti-correlated with the distribution of sensory information in cortical space. (A). Schematic illustration of spatial regulation: in each task condition, regions with higher alpha/beta power suppress local spiking, thereby reducing sensory encoding and information. (B). Sensory information (PEV for stimulus identity) carried by spiking activity at low- and high-alpha/beta electrodes in dlPFC (top) and vlPFC (bottom). Horizontal grey bars on the x-axis represented significant PEV differences between low- and high-alpha/beta power (p<0.05, two-sample t-test). Error bar: mean ± SE. Sample sizes (low/high alpha-beta): dlPFC spiking (n = 650/427), vlPFC spiking (n = 198/262). (C). Same as (B) but for sensory PEV carried by gamma at low- and high-alpha/beta electrodes in dlPFC (top) and vlPFC (bottom). Error bar: mean ± SE. Sample sizes (low/high alpha-beta): dlPFC gamma (n = 672/730), vlPFC gamma (n = 365/909). (D). Example spatial patterns from a single session in dlPFC (top) and vlPFC (bottom) showing time-averaged alpha/beta power, sensory PEV by gamma, and sensory PEV by spiking during the sample presentation period. Each pattern was normalized individually. (E). Correlation between spatial patterns of alpha/beta power and sensory PEV of gamma and spiking in dlPFC (top) and vlPFC (bottom). *** p<0.001, *: p<0.05, n.s.: not significant, one-sample t-test. Error bar: mean ± SE. Sample sizes: dlPFC (gamma: n = 22, spiking: n = 15), vlPFC (gamma: n = 20, spiking: n = 18). (F). Correlation between spatial patterns of alpha/beta power and sensory PEV of spiking for the same category (within-category) and different categories (between-category), for dlPFC (top) and vlPFC (bottom). *** p<0.001, *: p<0.05, n.s.: not significant, one-sample and two-sample t-test. Error bar: mean ± SE, n = 51 sessions.

First, we looked for evidence consistent with alpha/beta regulating where spiking sensory information is expressed. We analyzed data from the Single-image Task (**Figure 1D**), where multi-electrode Utah arrays were implanted in both dlPFC and vlPFC. We grouped LFPs/spiking by their positions in the electrode array. For each electrode, we computed the time-averaged alpha/beta power during sample presentation. We then split electrodes into two groups—those with higher or lower alpha/beta power across electrodes. Finally, we computed PEV reflecting sample object identity separately in spiking and gamma for each single electrode in order to construct spatial maps of sensory information (see **Methods**).

As predicted, spiking activity on electrodes with lower alpha/beta power exhibited significantly greater sensory information than spiking activity on electrodes with higher power (**Figure 5B**). This differentiation was significant but weaker in gamma (**Figure 5C**). To visualize these spatial relationships, **Figure 5D** showed example maps of the alpha/beta power, sensory information in gamma, and sensory information in spiking from a representative session. It showed anti-correlation between the spatial patterns of alpha/beta power and of sensory information in spiking. Across all sessions, we found a significant negative spatial correlation between alpha/beta power and sensory information in spiking (**Figure 5E**). This supports the idea that alpha/beta regulates the spatial distribution of sensory information in spiking.

Next, we assessed how this spatial anti-correlation shifted with changing task conditions, using data from the Category Task. Because alpha/beta patterns flexibly reorganize with category (**Figure S5E-H**), the spatial pattern of spiking sensory information should also reorganize accordingly. To examine this, we computed spatial correlations between alpha/beta power and spiking sensory PEV within the same category, and between different categories. Within the same category, spiking sensory PEV remained negatively correlated with the corresponding alpha/beta pattern (**Figure 5F**). By contrast, when comparing spiking sensory PEV from one category to alpha/beta patterns from the other, this anti-correlation weakened significantly and became more uncorrelated (**Figure 5F**). This suggested that the expression of spiking sensory information *remapped* with the spatial reorganization of alpha/beta power. It did so in a way that the alpha/beta pattern in one category better predicted the spatial distribution of spiking sensory information within the same category than in the other. Together, these results support Prediction 4.

### Prediction 5: Alpha/Beta Correlates with Behavior

If alpha/beta truly encodes task information in a way that influences the N s’ decisions, we would expect these signals also to differ between correct and incorrect trials and to correlate with trial-by-trial decisions.

We first tested this in the Recall version of Seq-image Task, where NHPs had to report the two sample items in the correct order. This design allowed us to isolate task-related errors. For many of the incorrect trials, NHPs chose the correct samples but in the wrong order, indicating an error of reversed order. Thus, we compared the neural information reflecting sample order between correct and order-error trials. We found that all signals carried significantly more order information in correct trials than in order-error trials (**Figure 6A**). This behavioral modulation was greatest for alpha/beta and spiking (**Figure 6B**). By contrast, information about sensory stimuli (i.e., sample identity) did not differ between correct and order-error trials for all signals (**Figure 6C** and **6D**). Similar results were also found for the Category Task by comparing correct vs. incorrect categorization trials (**Figure S6** and **S7**). This suggested that alpha/beta and spiking were crucial for maintaining the correct sequence order and category that influenced behavioral decisions.

**Figure 6.**
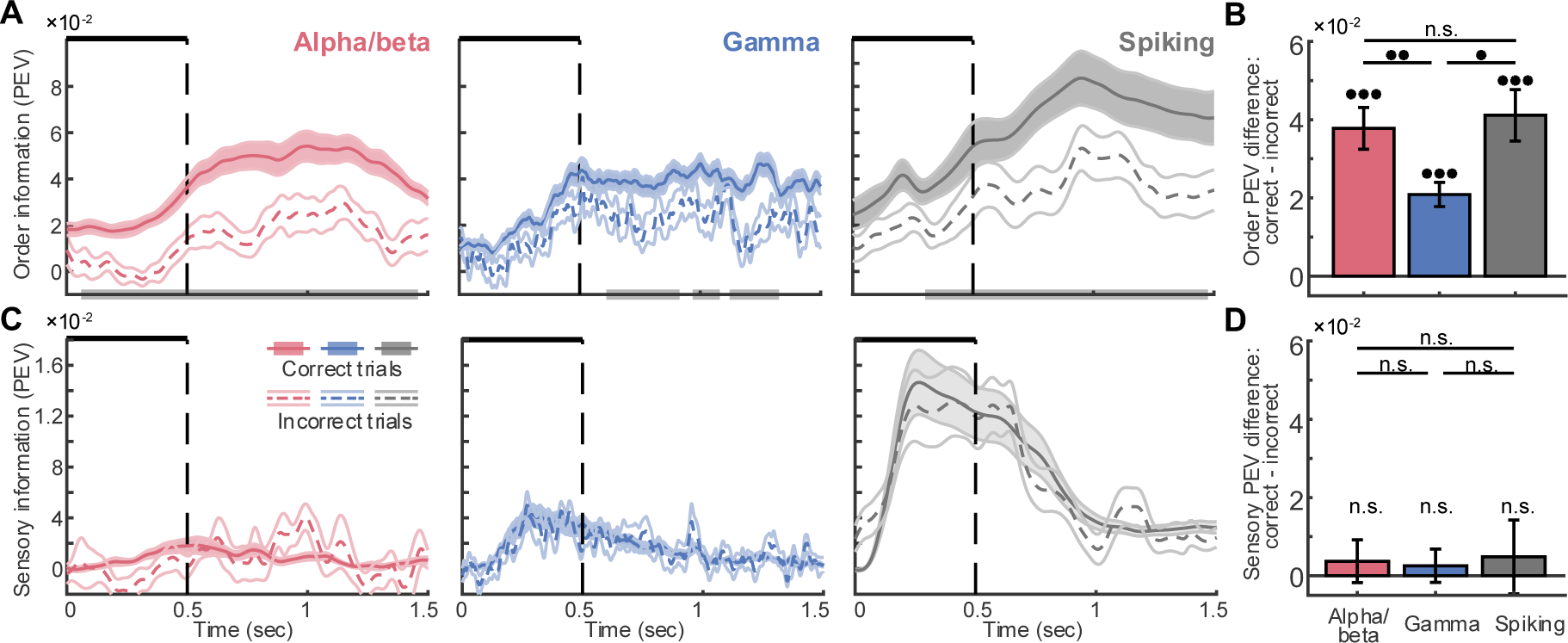
Alpha/beta and spiking reflect correct versus incorrect decisions about sequential order, but not about stimulus identity. (A). Task information (PEV) about sample order in Seq-Image task for correct trials (solid lines) and order-error trials (dashed lines), where NHPs chose the correct samples but in reversed order. Results are shown separately for alpha/beta (left), gamma (middle), and spiking (right). Horizontal grey bars on the x-axis indicate significant PEV differences between correct and error trials (p<0.05, two-sample t-test). Error bar: mean ± SE, n = 24 sessions. (B). Cumulative difference in order information (correct minus incorrect) across time for each signal type. *** p<0.001, **: p<0.01, *: p<0.05, n.s.: not significant, one-sample t-test for each signal type and Wilcoxon rank-sum test for across-signal comparisons. Error bar: mean ± SE, n = 24 sessions. (C). Same as in (A) but for sensory information about stimulus identity. (D). Same as in (B) but for sensory information.

We next turned to the Category Task to determine whether these signals correlate with NHP’s trial-by-trial decisions of category membership. We adapted a trial-by-trial analysis similar to Cohen & Maunsell, 2010 ^23^. Specifically, we projected each trial’s neural population data (time-averaged over sample presentation) onto an axis in population activity space that optimally separates the two learned categories on correct trials (see **Methods**). We then normalized projections onto this “category axis” so that a value of –1 corresponded to the average for Category-1 correct decisions and +1 to the average for Category-2 correct decisions. This provided a single-trial metric of population category coding strength. We then projected onto this axis the activity from the incorrect trials, which were not used to define the axis. If alpha/beta patterns are indeed relevant for categorization, we would expect the projections of the incorrect trials to be “shifted” toward the opposite, incorrect, category.

We found the predicted shift in alpha/beta for error trials in which the incorrect category was chosen. **Figure 7A** showed an example session in which alpha/beta projections for incorrect decisions on Category-1 trials shifted toward the Category-2 distribution, and vice versa. We quantified this effect by computing |Δ|, the absolute deviation between the mean projections of correct and incorrect category decisions (see **Methods**). Larger |Δ|values reflect a stronger shift in population activity on error trials, on average. We applied the same analysis to gamma and spiking (**Figure 7B**). We found the average |Δ|deviations across sessions differed significantly from zero (one-sample t-test, p<0.001) for all signals under both low- and high-abstractness conditions. However, alpha beta projections showed the greatest deviations |Δ|under high-abstractness conditions as compared to gamma and spiking, in both dlPFC and vlPFC.

**Figure 7.**
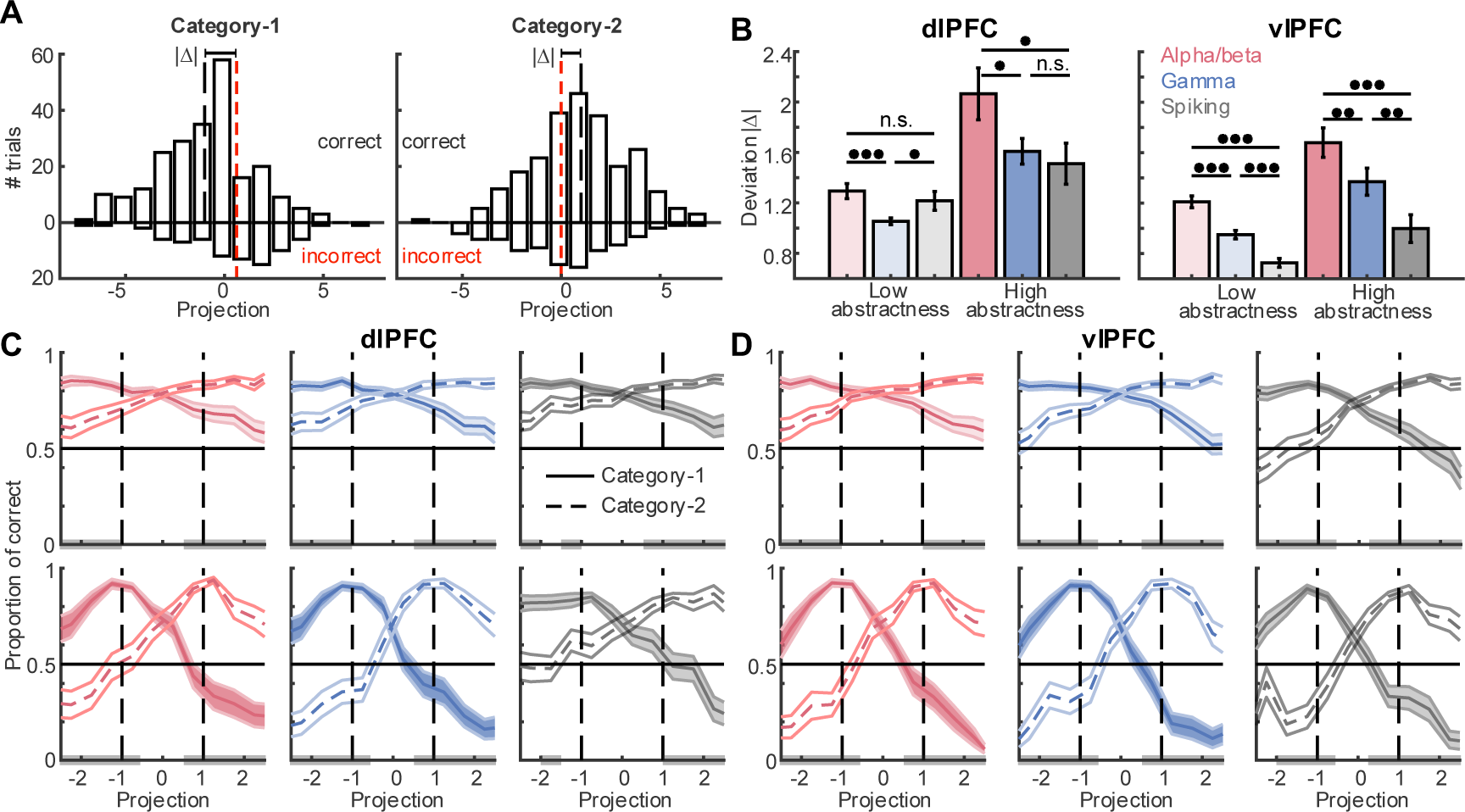
Alpha/beta patterns reflect category errors and predict trial-by-trial decisions. (A). Histogram of single-trial projections of dlPFC alpha/beta activity onto the category axis for correct (upward bars) and incorrect (downward bars) trials in an example session, pooled across low- and high-abstractness conditions. (B). Deviation (|Δ|) between mean projections of correct and incorrect trials under low- and high-abstractness conditions, shown for alpha/beta, gamma, and spiking in dlPFC (left) and vlPFC (right). *** p<0.001, n.s.: not significant, one-sample t-test for each signal type and Wilcoxon rank-sum test for across signals. Error bar: mean ± SE, n = 51. (C). Correct choice probability as a function of projection values along the category axis for alpha/beta (left), gamma (middle), and spiking (right) in dlPFC. Solid line: trials where Category-1 is the correct category; dashed line: Category-2 is the correct category. Grey bars on the x-axis represent significant differences between the choice probability of category-1 and category-2 (p<0.001, Wilcoxon rank-sum test). Error bar: mean ± SE, n = 51. (D). Same as (C) but for vlPFC.

To further characterize the direction of the shift, we analyzed the signed projections of incorrect trials. The shift toward the opposite category direction should reveal projections that consistently reversed sign relative to the correct category. We found this robust effect in alpha/beta projections across all conditions. In both dlPFC and vlPFC, for both low- and high-abstractness levels (**Figure S8**, left). Gamma also exhibited significant sign reversals, but only under high-abstractness conditions in both regions (**Figure S8**, middle). Spiking showed a similar pattern, but only in dlPFC and only for high-abstractness (**Figure S8**, right). These results suggested that while all signals reflect categorization errors to some extent, the categorization errors were most consistently and strongly associated with a systematic shift in alpha/beta toward patterns reflecting the incorrect category. This underscored the critical role of alpha/beta in representing task conditions used to guide decisions.

We also found a strong correlation with behavioral performance on a trial-by-trial basis. **Figure 7C** and **7D** show the proportion of correct category decisions as a function of the single-trial population activity projections. Trials with projections closer to the ideal value for the given category (*x*-axis values of –1 for Category-1, +1 for Category-2) resulted in the highest probabilities of correct decisions. Trials with projections toward the opposite end of the category axis (*x*-axis values of +1 for Category-1, –1 for Category-2) resulted in more incorrect decisions. In between these values, choice probability varied in a graded manner. The slopes of the choice-probability curves were similar across all signal types. This likely reflected the fact that, although alpha/beta showed a larger mean separation between correct and incorrect decisions, it also had greater trial-to-trial variability. In contrast, spiking and gamma showed smaller mean shifts but also lower variance (**Figure S9**), resulting in comparable overall predictive performance across signals.

All together, these results support Prediction 5. Alpha/beta patterns reflected sequence order errors. They reflected category errors. They reflected the N s’ trial-by-trial category decisions. We also observed that spiking and gamma signals correlated with behavioral performance. This is consistent with the theory: if alpha/beta oscillations regulate where sensory-related spiking can emerge, then behaviorally relevant information should also be reflected in spiking and associated gamma activity.

## Discussion

Our results provide evidence for the central predictions of Spatial Computing Theory across multiple cognitive tasks. Alpha/beta power conveyed minimal sensory information (i.e., the items held in WM) but robustly carried information about task conditions (i.e., task rules, order, and categories). Alpha/beta power carried stronger task information with greater cognitive demands. By contrast, neuronal spiking activity encoded both task and sensory information (**Predictions 1 and 2**). Alpha/beta power was spatially organized into patterns reflecting task conditions (**Prediction 3**). Sensory information carried by spiking was spatially anti-correlated with alpha/beta patterns. Recording sites with lower alpha/beta power had significantly stronger sensory information. This spatial anti-correlation suggested alpha beta’s role in spatially organizing information processing in the cortex (**Prediction 4)**. Lastly, alpha/beta power closely tracked N s’ behavioral performance. Alpha beta power reflected misattributions of the current task condition, whereas sensory information remained unaffected. Alpha beta patterns also correlated with N s’ trial-by-trial decisions about task conditions, especially under higher cognitive demands (**Prediction 5**). Collectively, these findings validate predictions of Spatial Computing Theory ^1^, demonstrating that alpha/beta oscillations organize cortical activity spatially to support context-dependent cognitive processing.

More broadly, our results are consistent with prior work showing that alpha/beta-band power and coherence in PFC reflects task rules and rule shifts ^24,25^. Our observations of increased context information in alpha/beta with increased cognitive demands parallels results from Wutz, et al., 2018 ^13^, who reported increased dlPFC alpha/beta power under similar conditions. Our findings go further by directly quantifying the amount of context information in alpha/beta and showing how it varies with task demands. Alpha/beta effects have also been observed across different task periods of simpler working memory tasks ^16,26–28^, putatively reflecting their differing requirements for task-related usage of information in working memory (i.e. encoding vs. maintenance vs. readout from working memory). These findings support the broader idea that low-frequency oscillations flexibly adapt to cognitive demands and help coordinate cortical processing.

In addition to these functional distinctions between signal types (alpha/beta vs gamma and spiking), we also observed overlaid broad differences between PFC subregions. We found stronger overall alpha/beta and stronger correlations with cognitive factors in dlPFC. We observed stronger spiking activity and stronger correlations with bottom-up sensory information in vlPFC. This is consistent with prior studies showing that dlPFC is more involved in maintaining abstract rules and goals, whereas vlPFC is more engaged in stimulus-driven encoding and maintenance of object features ^13,17–19^. It’s also consistent with the stronger connectivity between sensory cortex and vlPFC ^29,30^, and the stronger connectivity between dlPFC and areas involved in decision-making and motor output ^31,32^. Thus, these two areas may reflect a functional gradient, where top-down cognitive functions (and their associated signals) are more strongly represented in dlPFC and sensory-related functions (and their associated signals) are more strongly represented in vlPFC.

Previous studies have linked alpha/beta oscillations with behavioral performance ^33–35^, as we have in this study. We found that behavioral correlates in the alpha/beta signals were comparably strong to those observed in spiking activity. This suggests that the spatial patterns observed in alpha/beta oscillations are not merely reflective of internal state, but likely play an active role in shaping the neural processes that lead to behavior. They actively structure the neural landscape in ways that influence downstream decision-making.

Our findings align closely with conceptual advances from human EEG and MEG research. In humans, alpha oscillations have been widely implicated in the allocation of neural resources via the inhibition of task-irrelevant brain areas, particularly under top-down control ^36–38^. While most human studies focus on posterior cortical areas, modulations of frontal alpha activity have also been reported during goal-directed tasks, including language and working memory ^39,40^. Our data extend this framework by showing in non-human primates that alpha/beta oscillations in prefrontal cortex not only reflect top-down control but also shape where spiking-based sensory representations emerge. Our results further show that this control can be exerted in spatial patterns *within* a cortical area, and that these patterns show correlations with behavior. By linking alpha/beta patterns to both local excitability and behavioral performance, our results bridge a critical gap between non-human primate and human research. This suggests that low-frequency oscillations implement a shared control strategy across species and cortical areas.

Our results support the view that alpha/beta oscillations exert inhibitory control over local neural populations ^41,42^. Prior studies have shown that alpha/beta power decreases in cortical areas engaged by a task and increases in disengaged areas ^43–45^. In macaque cortex, increased alpha/beta power has been associated with reduced local spiking activity, suggesting a suppressive gating function ^28,35,46–48^. Consistent with this, we observed a strong inverse relationship between alpha/beta power and content-related spiking. This correlational relationship supports an inhibitory role for alpha/beta. Future work, including causal analyses and modeling, will be needed to clarify the directionality of this relationship and provide a mechanistic account of how alpha/beta might regulate spiking activity.

One possible neural circuit mechanism underlying this spatial regulation involves local inhibitory interneurons. Recent evidence, both experimental ^49–52^ and computational ^53–55^, suggests that alpha/beta oscillations may be generated by somatostatin-expressing (SST) interneurons. These interneurons are well-positioned to mediate top-down modulation ^56,57^. Top-down inputs could locally recruit SST cells, leading to an increase in the local alpha/beta power and targeted suppression of nearby excitatory neurons and spiking. In this way, the spatial structure of alpha/beta activity could emerge from selective activation of inhibitory circuits, which in turn sculpt where content-related spiking is permitted or suppressed.

Our findings build on prior work showing that working memory is implemented through interactions between alpha/beta oscillations and brief spiking/gamma bursts ^16,28^. More recently, Lundqvist et al. (2023) ^1^ formalized these ideas into Spatial Computing theory, proposing that cortical space is used as a computational dimension for cognitive control. They showed that low-dimensional oscillatory activity carries control-related signals. While that study established key principles of Spatial Computing, several important questions remained open. This includes how spatial patterns of neural activity reorganize with context and scale with cognitive demand, and whether these dynamics relate to behavior. Our results extend this framework by providing direct empirical tests of these predictions. Our results suggest that alpha/beta oscillations not only gate information in time but also sculpt the spatial landscape of cortical activity, determining where spiking-based sensory representations emerge. This offers strong support for the idea that cortical space functions as an active computational resource, with alpha/beta rhythms regulating both the location and timing of information expression.

This spatial regulation offers a mechanistic basis for how the brain flexibly reconfigures its representational geometry with task demands. This is closely aligned with theories of subspace coding ^3,4^ and nonlinear mixed selectivity ^5–8^. Subspace coding proposes that different task contexts activate orthogonal patterns of population activity, minimizing interference between representations. Mixed selectivity refers to neurons that respond to specific combinations of stimuli and task context. It is essential for flexible cognition^6^. While these properties have been observed across the brain, the underlying mechanism has remained unclear. Our results suggest that alpha/beta oscillations provide that mechanism by dynamically regulating where in cortical space certain subspaces and selective combinations can emerge. By spatially suppressing spiking activity, alpha/beta patterns may reduce the overlap between neuronal populations recruited in different contexts, leading to more orthogonal population subspaces. This selective recruitment could also support the emergence of nonlinear mixed selectivity, as neurons participate in different combinations depending on where alpha/beta inhibition is released. In doing so, Spatial Computing explains how a single cortical circuit can express a wide range of task-dependent representations without structural rewiring.

Gamma is typically associated with bottom-up sensory processing and closely tied to local spiking activity ^58–60^. We found gamma carried some sensory information, though to a much lesser extent than spiking. This likely reflects its lower spatial and temporal resolution relative to spiking activity. Gamma also carried some task information and was correlated with behavioral choices. These findings suggest that while gamma is not traditionally considered a carrier of top-down signals, it may inherit context-related structure through its close coupling with spiking activity that is spatially regulated by alpha/beta oscillations. In this sense, gamma may act as an intermediary, reflecting both the bottom-up content and the spatial constraints imposed by top-down control. Prior work has shown that gamma synchronization is modulated by attention and correlates with behavioral measures, including perceptual accuracy and reaction time ^61,62^.

Taken together, our results highlight Spatial Computing as a framework for understanding how the brain flexibly controls cognitive processes. By showing that alpha/beta oscillations dynamically shape when and where content can be expressed, we report a spatial mechanism for cognitive control. This shifts the focus from fixed anatomical circuits to dynamic, spatially structured population activity as the substrate of cognition. Spatial Computing offers a unifying principle that explains how diverse cognitive operations, such as working memory, rule application, categorization, and decision-making, can emerge from a shared circuitry guided by oscillatory control. More broadly, it suggests that cortical space itself is a computational dimension, repurposed in real time to serve changing behavioral goals. This view opens new avenues for understanding the flexible and adaptive nature of neural computation, and for building models of cognition grounded in the analog dynamics of brain activity.

## Acknowledgments

We thank Jordan DeFarias and Meredith Mahnke for technical assistance in data collection. We thank all members of the Miller Lab for helpful comments and discussions.

## Funding

This study was funded by Office of Naval Research W911NF2410228 (E.K.M.), Office of Naval Research MURI N00014-23-1-2768 (E.K.M.), Freedom Together Foundation (E.K.M.), The Picower Institute for Learning and Memory (E.K.M.), and ERC starting grant Hot-Coal WM 949131 (M.L.). This material is based upon work supported in part by ARO award W911NF-24-1-022 (E.K.M.).

## Author Contributions

M.R.W, R.F.L., S.L.B., and M.L. collected the data. S.L.B. curated the data. M.L. originated the concept of spatial computing. Z.C. and S.L.B. formalized the central predictions of spatial computing. Z.C. analyzed the data to test the predictions. Z.C., S.L.B., and E.K.M. wrote the manuscript.

## Competing interests

The authors declare that they have no competing interests.

## Data and materials availability

All data needed to evaluate the conclusions in the paper are available upon reasonable request from the Corresponding Author.

## Methods

### Non-human primate (NHP) subjects

We analyzed data from three previous studies. The Seq-image Task was from ^9,14^, the Single-image Task was from ^1,10,11^, and the Category Task was from ^12,13,63–65^. For details of experimental design and data collection, please see those publications.

Briefly, each study involved two Rhesus macaques (*Macaca mulatta*) that were trained on a behavioral task using positive rewards (juice), until their performance indicated a solid understanding of all the task requirements. The Seq-image Task included NHPs S (female, 6 years, 6.5 kg) and A (male, 4 years, 10 kg). The Single-image Task included NHPs T (male, 17 years, 13 kg) and D (male, 8 years, 10 kg). The Category Task included NHPs P (female, 8 years, 9 kg) and G (male, 9 years, 13 kg). All NHPs were experimentally naive prior to the experiments analyzed here, except for NHP T, who had previously participated in studies of categorization ^63^ and working memory ^65^; NHP P, who had participated in a previous study of category learning ^66^; and NHP G, who had participated in a study of rule-based categorization ^64^. NHPs were pair-housed, on a 12-hr day/night cycle, and in a temperature-controlled environment (80° F). All procedures followed the guidelines of the Massachusetts Institute of Technology Committee on Animal Care and the National Institutes of Health.

### Datasets analyzed

For the Seq-image Task, NHPs were trained on two versions of a sequence working memory task. Blocks of each task version—Recognition and Recall—were interleaved, 100–250 trials per block. In both versions, a sequence of two sample images was presented (500 ms each, 1 s blank delay between them). The identity and order of the images then had to be maintained in working memory over a brief delay (1 s). In the Recognition version, a test sequence of two images was then presented, which required a bar-release response during the presentation of the second image if the test images matched the samples in both identity and order. Otherwise, subjects withheld their response until a subsequent matching sequence was shown. In the Recall version, an array of three test images was shown simultaneously, and the subject was required to saccade to the two images that matched the samples in the same order as initially presented. There was no explicit cue to indicate that the task had switched, as it was obvious from context during the first trial after the block switched. NHPs typically performed 2–4 blocks of each task during each recording session. Performance was well above chance on both versions of the task. An infrared-based eye-tracking system (ISCAN) continuously monitored eye position at 60 kHz. In each session, eight tungsten microelectrodes (FHC) were lowered through custom grids using custom-made independently moveable microdrives (Crist). LFPs were referenced to ground, band-pass filtered, and recorded at 1 kHz (Plexon Multichannel Acquisition Processor). Spikes were detected from high-frequency data using an online manually-set amplitude threshold, and manually sorted into isolated single units offline (Offline Sorter, Plexon). For NHP S, a DC-coupled headstage, LFP filtering from 3.3–88 Hz, and spike filtering from 154–8800 Hz was used. For NHP A, an AC-coupled headstage, LFP filtering from 0.7–170 Hz, and spike filtering from 250–8000 Hz was used.

For the Single-image Task, NHPs were trained on a delayed match-to-sample WM task. Each trial began with fixation for 500 ms, followed by a sample object, blank memory delay, and then a free-viewing two-choice test. During the test period, NHPs were required to saccade to the remembered sample and hold fixation there. An infrared-based eye-tracking system (Eyelink 1000 Plus, SR-Research, Ontario, CA) continuously monitored eye position at 1 kHz. Neural signals were recorded with four chronically implanted 64-electrode (8×8) “Utah” arrays (1.0 mm length, μm spacing, iridium-oxide tips; Blackrock Microsystems, Salt Lake City, UT). Arrays were implanted in both hemispheres of dorsolateral PFC and ventrolateral PFC. LFPs were referenced to low-impedance subdural reference wires, AC-coupled, band-pass filtered (0.05–300 Hz), and recorded at 1 kHz (Blackrock Cerebus). Spikes were detected from high-frequency data (250–5000 Hz) using a manually-set amplitude threshold and spike-sorted online using time-amplitude windows.

For the Category Task, NHPs were trained to categorize two categories of dot-pattern category exemplar stimuli, each of which was defined by random distortions around a prototype. Each prototype consisted of 7 pseudo-randomly placed dots. The exemplars were selected by randomly sampling locations from a normal distribution centered at each prototype dot location. The session was split into blocks of trials, with exponentially increasing numbers of category exemplars and increasing levels of distortion around each prototype in successive blocks. The first block featured two exemplars of each category, and each subsequent block doubled the number of exemplars, cumulatively keeping all exemplars from previous blocks and adding an equal number of new exemplars. Each block, the variance of the normal distributions around the prototype dots, which was used to generate each exemplar (i.e. the distortion), progressively increased. NHPs moved from one block to the next when a learning criterion was achieved. Neural signals were recorded with two chronically implanted 64-electrode (8×8) “Utah” arrays (1.0 mm length, μm spacing, iridium-oxide tips; Blackrock Microsystems, Salt Lake City, UT). Arrays were implanted unilaterally in dorsolateral PFC and ventrolateral PFC. LFPs were referenced to ground, AC-coupled, band-pass filtered (0.75–300 Hz), and recorded at 1 kHz (Blackrock Cerebus). Multi-unit spiking activity was obtained by offline high-pass filtering and thresholding of the 30 kHz-sampled raw broadband signals. A channel was defined as recording spikes (multi-unit (MUA) channel) when its spike rate averaged over trials and sample presentation period (0–1 sec) was larger than 1 Hz.

Note that any global fluctuations that might have been introduced via the reference wire (Single-Image Task) or ground (Seq-Image Task, Category Task) would be expected to, if anything, work against our primary results. Our results rely on the fact that alpha/beta oscillations exhibit *specific spatial patterns*. Since global fluctuations are, by definition, consistent across all electrodes, they would actually mask any such patterns. Further, to the extent that global fluctuations reflect noise or other non-task-related factors, they would also work against our results showing that alpha/beta patterns differ between task conditions.

### Signal processing

In all datasets, local field potentials (LFPs) were first preprocessed by removing the DC offset, and removing the 60 Hz line noise and its harmonics using an adaptive sinusoid fit method. Subsequently, signals were band-pass filtered between 0.5 and 300 Hz.

For spectral analyses of the LFPs, we applied Morlet wavelet analysis ^67^ after subtracting the evoked potentials phase-locked to trial events, estimated using the grand mean signal across all trials. The Morlet wavelets had a fixed width of 6 cycles, with center frequencies ranging from 2^1^ to 2^7.5^ Hz in increments of 0.2 (on a logarithmic scale). Resulting spectrograms were then normalized to correct for the ∼1/frequency baseline distribution of power by multiplying power at each frequency by the frequency itself ^68,69^. To obtain band-specific oscillatory power, we averaged the spectral data within the alpha/beta (10–30 Hz) and gamma (30–150 Hz) frequency bands. Note that, without the above correction for the 1/frequency distribution of power, these band-pooled power estimates would be heavily biased toward the lower-frequency end of each band. Oscillatory power was calculated with a temporal resolution of 1 ms.

Neural spiking data across all datasets were smoothed using a Gaussian kernel with a width of 50 ms. The smoothed spiking activity is referred to as the spike rate, with a temporal resolution of 1 ms. Both LFP band power and spike rate data were baseline-corrected by subtracting the mean activity (averaged across trials and time points) in the time interval preceding stimulus onset: [–1, 0] s relative to sample onset for Seq-image Task and Category Task and [–0.5, 0] s for Single-image Task.

In addition to these standardized preprocessing steps, dataset-specific procedures from the original studies were applied as necessary; further details are provided in those publications. Briefly, for the Seq-image Task, sessions containing high-power, broadband frequency artifacts were excluded from analyses. In the Single-image Task, channels identified as having poor electrode contact or significantly elevated noise amplitudes, based on visual inspection, were removed prior to analysis. Unless specified otherwise, all computations below were performed on a session-by-session basis.

### Information quantification using PEV

To quantify the amount of information carried by each neural signal type (alpha/beta power, gamma power, and spike rate), we computed the bias-corrected percentage of explained variance (PEV) ^70^ across trials under different task conditions. For different analyses, this quantification was performed at both the population level and the level of individual electrodes or units.

At the population level, we first defined an optimal one-dimensional readout axis based on all correct trials. At each time point, neural population activity (LFP power or spike rate) formed distinct clusters in a high-dimensional space, where each dimension represented one electrode or unit. Each data point corresponded to the neural response from a single trial, and clusters represented two distinct experimental conditions (i.e., a binary classification). For task information, these conditions included Category-1 vs. Category-2 (Category task), first vs. second presentation order (Seq-image task), or Recognition vs. Recall rule versions (Seq-image task). For order information in the Seq-image Task, we held the image shown at both sequence orders constant to reduce the effect of stimulus identity on the variance. Each image shown gave us two separate clusters corresponding to first and second sequence order. For sensory information (e.g., stimulus identity), each stimulus pair formed two separate clusters.

For each binary contrast, we computed the linear discriminant axis, which reflects the optimal linear decoder that maximizes the separation between classes ^71,72^ and the Fisher information ^73^. The optimal axis was calculated as the product of the pooled inverse covariance matrix (𝛴) of the two clusters and the vector connecting their cluster means: 𝛴^−^*^1^*(𝜇*_1_*− 𝜇*_2_*). After averaging this optimal axis across time (and across stimulus pairs for stimulus identity, or across the images shown for order), we projected the original high-dimensional data onto this one-dimensional axis. This step effectively reduced the dimensionality of the neural data, facilitating the computation of PEV analogous to analyses conducted at the single electrode/unit level.

For the resultant one-dimensional projected population data, as well as for single electrode/unit data, we applied a one-way ANOVA, grouping trials according to different experimental conditions (e.g., stimulus identity, task categories, task rules). Only correct trials were included. A bias correction was applied to PEV estimation (𝜔*^2^* formula) to address potential biases associated with small sample sizes ^74^:

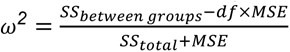

where 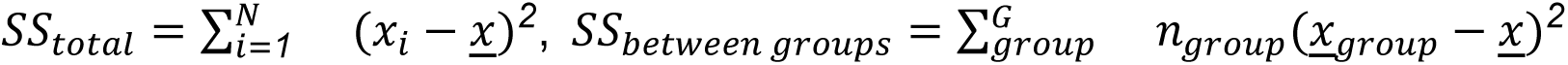, df is the degrees of freedom (i.e., the number of groups − 1) and M is the mean squared error 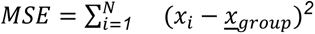. Consequently, bias-corrected PEV 𝜔*^2^* provided a robust measure of the amount of information encoded by modulations in neural spiking rates or oscillatory power at both the population and single-electrode/unit levels.

For the Seq-image Task, the sensory PEV was computed based on the trials of the Recognition and Recall versions separately for each session. They were aggregated across all the sessions of the two NHPs. The task PEV was computed by all the trials together with the two groups given by “ecognition” and “ecall”. For the order information, we separated each trial into two temporal segments: sample 1 + delay and sample 2 + delay. These two segments from each trial gave the two groups for PEV. The order PEV was computed for Recall trials only so it could be compared with the PEV of order-error trials, which are most easily identifiable in the Recall version.

For the Category Task, the sensory PEV was computed only over trials from the first block, where the number of exemplar dot patterns was small. This minimized the effects of category learning on sensory representation. The category information was computed for the blocks later than block three, by which NHPs typically learned the category ^12^.

To estimate the information during error trials, we projected incorrect single-trial neural responses onto the optimal one-dimensional axis derived from the correct trials. The same one-way ANOVA procedure described above was subsequently applied to these projections.

### Low- and high-abstractness based on exemplar distortion

For the Category Task, we classified exemplars into low- and high-abstractness groups using the same criterion as described previously by ^13^. Specifically, the dataset from each recording session was divided into two abstractness levels based on a critical distortion threshold of 1.1 degrees visual angle (DVA), corresponding to the inflection point in behavioral performance. Further methodological details are provided in ^13^. Analysis based on abstractness levels, for example, category PEV, was performed on the trials of low- or high-abstractness separately. For example, for the category information under low- and high-abstractness conditions, we computed the one-dimensional axis from the low- and high-abstractness trials separately and projected the data separately.

### Spatial clustering analysis of alpha/beta patterns

To characterize the spatial clustering and organization of alpha/beta power, we computed Moran’s I, a commonly used statistic to quantify the overall strength of spatial autocorrelation ^20–22^. It is essentially a distance-weighted correlation between all pairs of electrodes. As with other correlation measures, its value ranges from –1 (i.e., maximally spatially dispersed/anti-correlated) to +1 (i.e., maximally spatially clustered/autocorrelated). A value of 0 indicates a lack of correlation. We computed it separately on the baseline-subtracted alpha/beta power and on a measure of alpha/beta power selectivity (signed difference in power between task conditions). Note that per-electrode baseline subtraction removes any global power gradients ^27^, so any observed effects are unlikely to be explained by nonspecific spatial structure.

Moran’s I is defined as

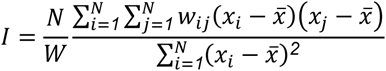

where 𝑁 is the number of electrodes indexed by 𝑖 and 𝑗, 𝑥_𝑖_ is the observed alpha/beta power (or selectivity) at the 𝑖-th electrode, 𝑥̅ is the averaged power (or selectivity) across all electrodes, 𝑤_𝑖𝑗_ are the elements of a matrix of spatial weights with zeroes on the diagonal, and 𝑊 is the sum of all *w*_*ij*_ (i.e., 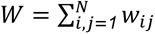). The spatial weights we chose were binary weights where nearest neighbors (i.e., the electrodes at a minimal distance from a given electrode) were set to 1 and all other weights were set to 0. We computed Moran’s I for the actual observed alpha beta patterns and patterns spatially shuffled across each recording array. As an alternative measure of spatial clustering, we also computed the absolute differences of baseline-subtracted alpha/beta power between nearest-neighbor electrodes, and also compared this to values for spatially-shuffled data (**Figure S5**). Both metrics gave consistent results.

### Spatial correlation between alpha/beta power and sensory PEV maps

To quantify the spatial relationship between alpha/beta power and the sensory information (PEV) encoded by gamma power and spiking, we first constructed spatial maps representing both power and sensory PEV.

For alpha/beta power, we first subtracted the baseline (pre-stimulus) values as described above. Then, for each electrode, power was averaged across all correct trials and across a specific time window during stimulus presentation (approximately 0.1–0.4 s for the Single-image Task and 0.2–0.6 s for the Category Task). This resulted in a vector of 64 alpha/beta power values recorded in each session. These values were then mapped onto the 8-by-8 electrode grid locations to build the power map.

For the sensory PEV maps, we first computed the PEV of stimulus identity separately for each electrode/unit at each time point. Note that, unlike the population PEV described above, this was computed separately for each individual electrode/unit, without any projection onto a population coding axis. We then averaged the resulting PEV time series within the same time window used for alpha/beta power averaging, producing a sensory PEV vector with dimension equal to the number of electrodes/units. For gamma activity, the resulting PEV vector was mapped directly to the grid locations of each electrode. For spiking activity, the approach varied slightly: for unsorted multi-unit activity (MUA) recorded in the Category Task, electrode locations were directly mapped. However, for sorted single-unit activity recorded in the Single-image Task, electrodes could have multiple units or none. When multiple units were recorded on the same electrode, their PEV values were averaged to yield a single electrode-level PEV value. Electrodes with no recorded units were excluded from spatial correlation computations.

Spatial correlations were computed using earson’s correlation coefficient. For each recording session, we correlated the alpha/beta power vector with the corresponding PEV vector, including only electrode locations containing valid data points (i.e., excluding electrodes with missing values because they recorded no units). Note that the observed anti-correlation between alpha/beta power and sensory PEV is unlikely to be attributed to variations in the 1/f spectral slope between trial epochs or experimental conditions ^15,75^. First, power spectra were corrected for 1/f scaling by multiplying power at each frequency by its corresponding frequency prior to any further analysis. Second, all spatial correlation analyses were performed within the same trial epoch and experimental condition (e.g., for a single memorized object identity), ensuring that any across-epoch or across-condition differences in 1/f slope could not account for the reported effects.

### Within- and between-category spatial correlation of alpha/beta power maps

For the Category Task, we assessed spatial correlations of alpha/beta power maps both within and between categories. Trials from each recording session were segmented into non-overlapping windows of 50 trials. Within each set, we computed spatial correlations separately and subsequently averaged the results across the trial windows. For each set of 50 trials, we constructed alpha/beta power maps by separately averaging trials from Category-1 and Category-2, both across trials and across the stimulus presentation temporal window. The between-category correlation was calculated directly between these two resulting spatial maps. For within-category correlations, trials from each category were randomly partitioned into two halves, and correlations were computed between spatial maps averaged from these halves. This random partitioning procedure was repeated 20 times, and the average correlation value was taken as the within-category correlation estimate. Finally, spatial correlation values obtained from the trial windows after the third block were averaged to yield the overall within- and between-category correlation estimates for each recording session.

### Single-trial behavioral performance analysis

Single-trial behavioral performance analyses were conducted following the procedure described by ^23^. For each recording session, neural responses (LFP power or spike rate) were first averaged across the stimulus presentation window for each individual trial, resulting in trial-specific population response vectors. Using only correct trials, we then calculated an optimal one-dimensional discriminant axis by multiplying the inverse covariance matrix of two response clusters (corresponding to Category-1 and Category-2) with the vector connecting their cluster means. Single-trial population responses were projected onto this discriminant axis, and the resultant scalar projections were normalized so that projections of -1 and +1 corresponded to the mean responses of correct Category-1 and Category-2 trials, respectively.

Note that we used all correct trials to estimate the decision axis to obtain a more stable and robust estimate. This was particularly critical when the data were further subdivided by category and abstractness level. This could, in principle, bias the axis toward better separation of correct trials than of error trials. However, this cannot account for the systematic shift of error trials to the *opposite* side of the axis relative to their correct category (**Figure S8**). Thus, our results are unlikely to be explained by overfitting to correct trials, but is consistent with genuine category mislabeling in error trials.

Incorrect trials exhibited a projection distribution shifted toward the opposite category: incorrect Category-1 trials shifted toward +1, while incorrect Category-2 trials shifted toward -1. For each session, we used the one-sided t-test to see whether this shift in the incorrect distribution was significant (p<0.05) or not. For those sessions with significant shifts, we quantified these shifts by computing the mean projection values separately for incorrect Category-1 and Category-2 distribution and then calculated the absolute differences (denoted as |Δ|) between these incorrect-trial means and their respective correct means (-1 for Category-1, +1 for Category-2).

To estimate the probability of correct choices as a function of scalar projection, we binned the projection values from all trials within each recording session and counted the number of correct and incorrect trials per bin. Given that the projections were normalized for each session, bin alignments were consistent across sessions, with -1 representing the average correct Category-1 response and +1 the average correct Category-2 response. Finally, we aggregated these binned trial counts across all sessions and both NHPs.

### Statistical methods

Statistical tests for significance were performed with a two-sided Wilcoxon rank-sum test (ranksum function in MATLAB) when samples were independent and with a two-sided t-test (ttest function in MATLAB) for paired samples or single samples. Correlations between two variables were computed as the Pearson correlation. Statistical significance was defined by a p value <0.05. The statistical details (correlation coefficient, p value, sample size n) are provided in the figures, figure legends, or the text of the Results section. The specific meaning of the sample size n is clarified when used.

---

**Figure S1.**
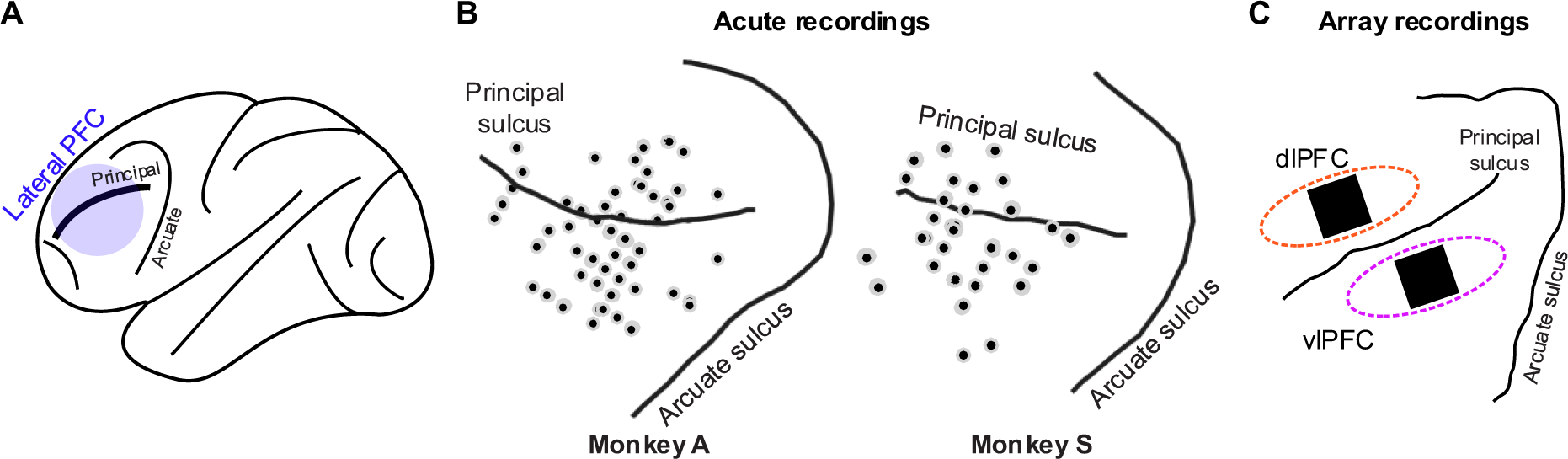
Recording sites in the lateral PFC across tasks. (A). A brain schematic showing the location of lateral prefrontal cortex (PFC) where all the recordings were performed. (B). Anatomical locations of recording sites for the Seq-image Task, for both NHPs. (C). Utah array locations in the dlPFC and vlPFC for the Single-image Task and Category Task.

**Figure S2.**
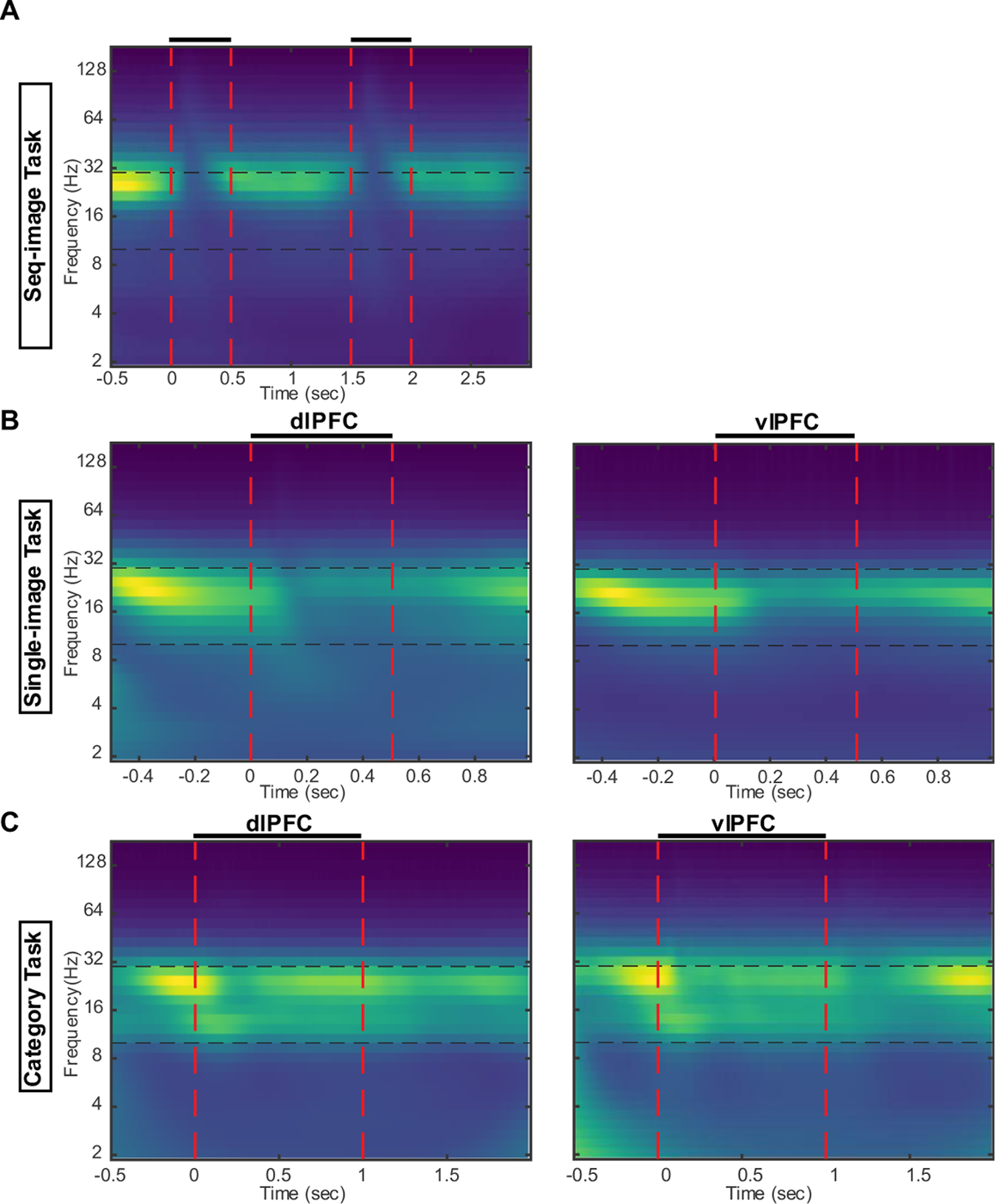
Trial-averaged power spectrograms for each dataset show band-limited alpha/beta oscillations. (A). Trial-averaged power spectrogram for the Seq-image Task. The alpha/beta band is in-between the black dashed lines. The red dashed lines indicate sample presentation. (B). Same as (A) but for Single-image Task. (C). Same as (A) but for Category Task.

**Figure S3.**
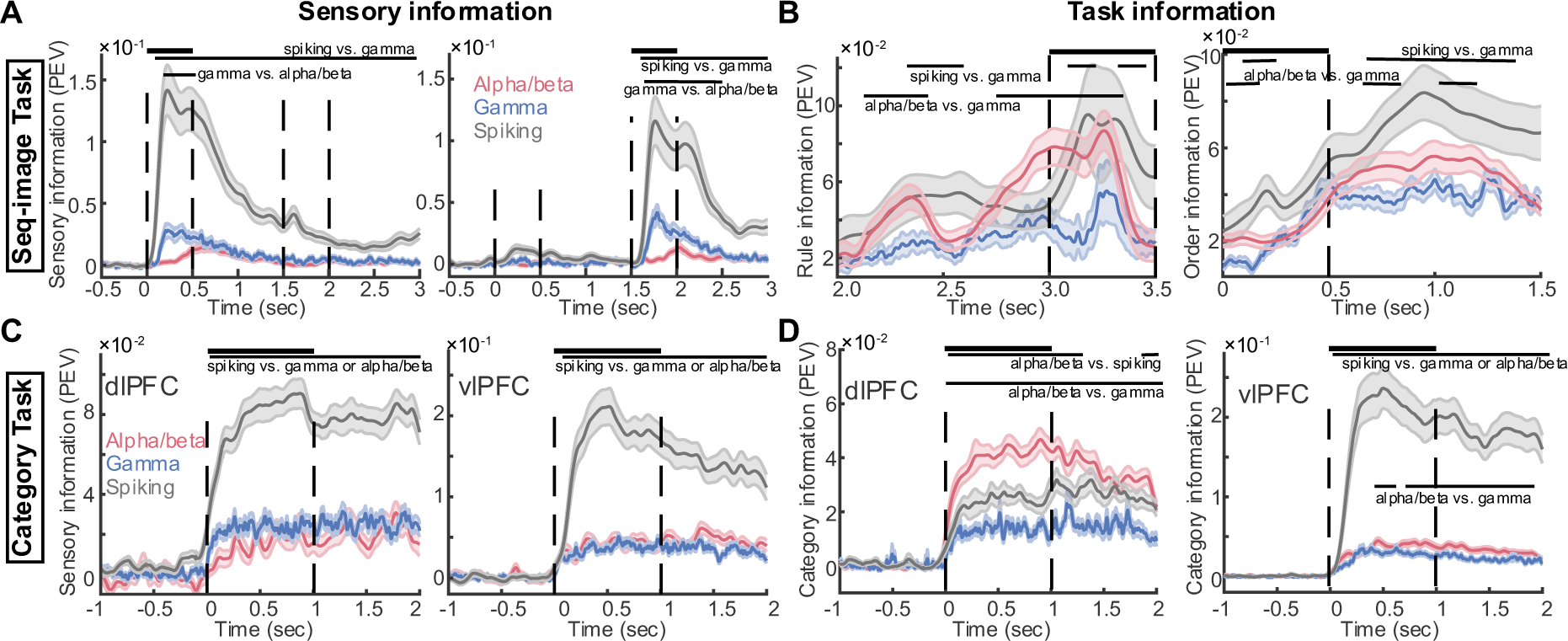
Paired significance test for sensory and task information. (A-B). Sensory (A) and task (B) information by each signal type in the Seq-image Task. Error bar: mean ± SE, n = 2×24 sessions for (A) and n = 24 sessions for (B). Horizontal black lines on top represent significant PEV values for pairwise comparison between signal types (p<0.001, Wilcoxon rank-sum). (C- D). Sensory (C) and task (D) information by each signal type in the Category Task. Left: PEV for dlPFC. Right: PEV for vlPFC. Error bar: mean ± SE, n = 51 sessions. Horizontal black lines on top represent significant PEV values for pairwise comparison between signal types (p<0.001, Wilcoxon rank-sum).

**Figure S4.**
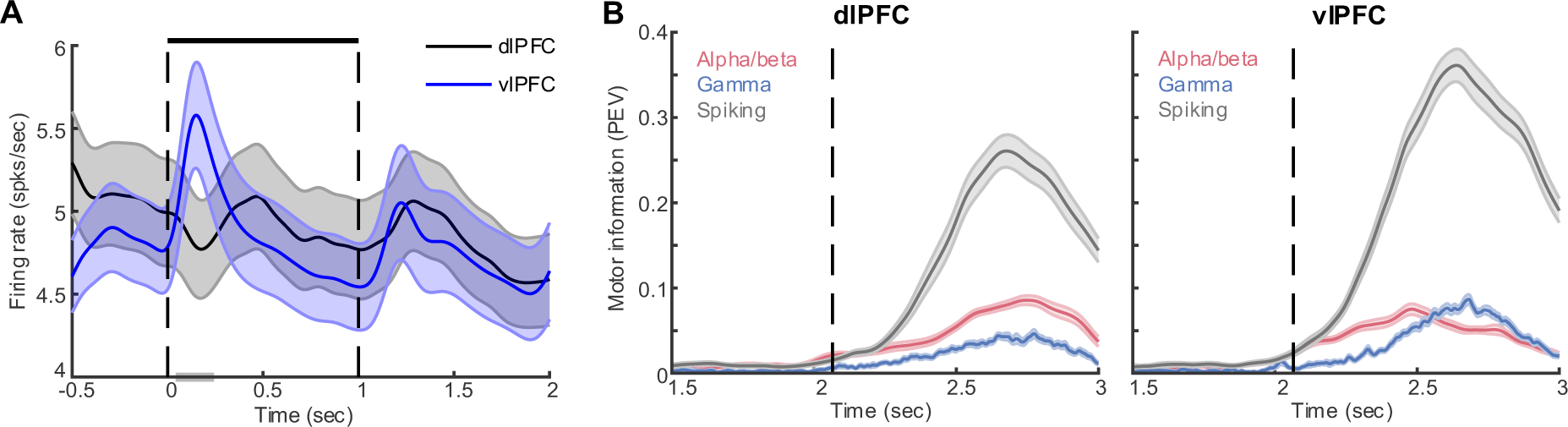
Firing rate and motor information for Category Task. (A). Firing rate (spikes per second) averaged over all dlPFC and vlPFC MUA channels (MUA channel = average spike rate between 0 and 1 s > 1). (B). Information quantified by PEV of the left vs. right saccade for dlPFC (left) and vlPFC (right). The vertical dash line indicates the time of test onset. Error bar: mean ± SE, n = 51 sessions.

**Figure S5.**
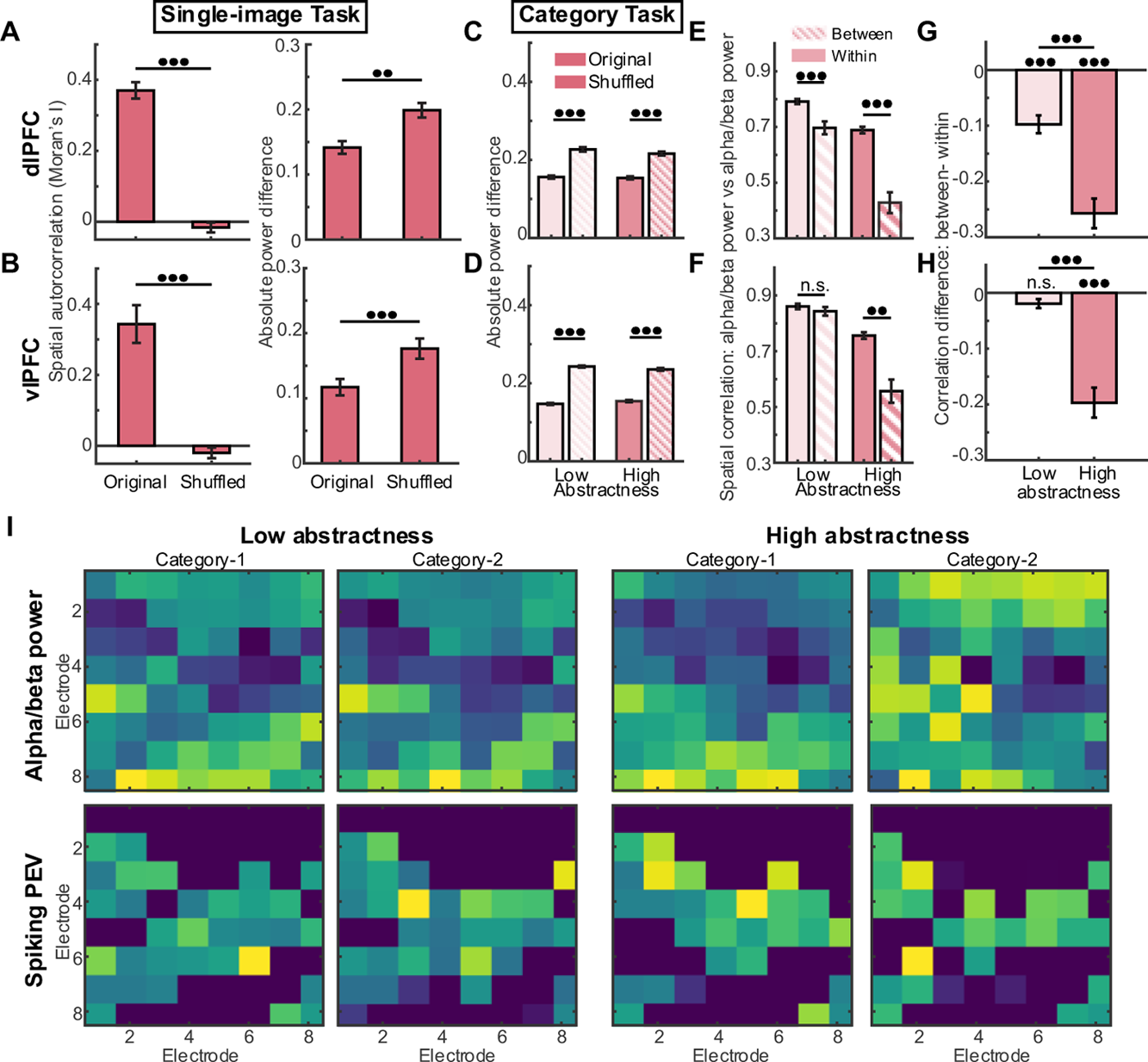
Spatial regulation of sensory information by alpha/beta patterns. (A). The spatial clustering of alpha beta power measured by Moran’s I (left) and absolute power difference (right) in dlPFC recorded from Single-image Task. The values are compared with randomly shuffled patterns (bar with stripes). *** p<0.001, **: p<0.01, two-sample t-test. Error bar: mean ± SE. Sample sizes: n = 22. (B). Same as (A) but for vlPFC of Single-image Task. Error bar: mean ± SE. Sample sizes: n = 20. (C). The spatial clustering of alpha/beta power measured by absolute power difference in dlPFC recorded from Category Task. *** p<0.001, **: p<0.01, two-sample t-test. Error bar: mean ± SE. Sample sizes: n = 51. (D). Same as (C) but for vlPFC of Category Task. (E). Spatial correlation of alpha/beta power computed for the same category (within) and for different categories (between) in dlPFC. *** p<0.001, **: p<0.01, two-sample t-test. Error bar: mean ± SE. Sample sizes: n = 51. (F). Same as (E) but for vlPFC. (G). Spatial correlation differences (between- minus within-category) for high- and low- abstractness conditions in dlPFC. *** p<0.001, **: p<0.01, n.s.: not significant, one- and two-sample t-test. Error bar: mean ± SE, n = 51. (H). Same as (G) but for vlPFC. (I). Example alpha/beta power maps with the corresponding sensory PEV carried by spiking for both categories under low and high- abstractness conditions.

**Figure S6.**
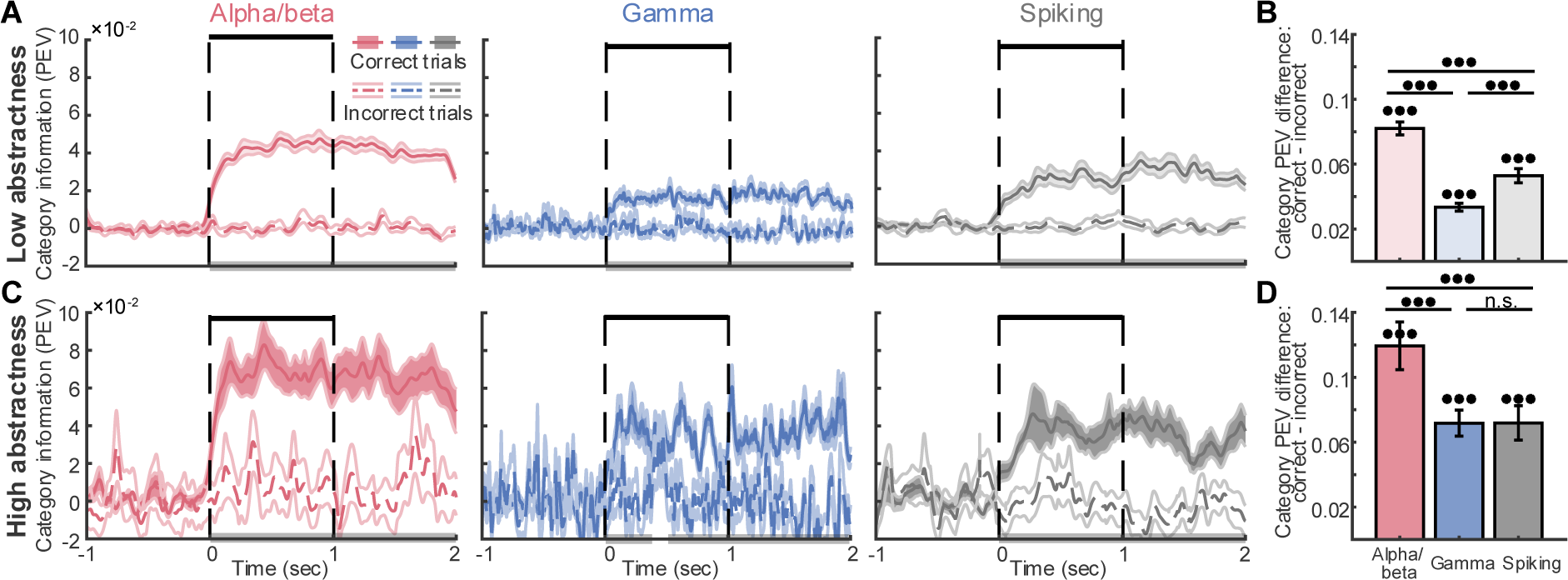
Category information (PEV) in dlPFC for correct and incorrect trials. (A). Category information (PEV) for correct trials and incorrect trials for alpha/beta (left), gamma (middle), and spiking (right) under low-abstractness. Horizontal grey bars on the x-axis represented significant PEV differences between correct- and incorrect conditions (p<0.05, two-sample t-test). Error bar: mean ± SE, n = 51. (B). Cumulative difference in category information (correct minus incorrect) across time for each signal type. *** p<0.001, n.s.: not significant, one-sample t-test for each signal type and two-sample t-test for across signals. Error bar: mean ± SE, n = 51. (C-D). Same as (A-B) but under high-abstractness.

**Figure S7.**
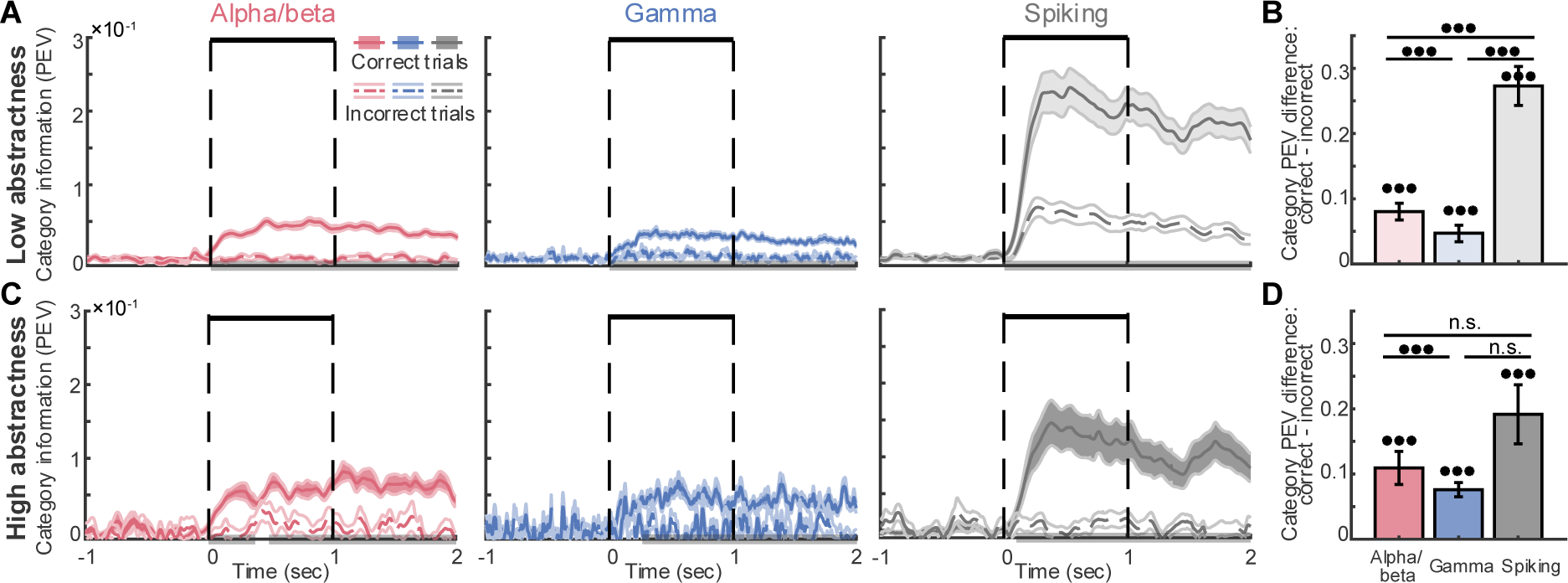
Category information (PEV) in vlPFC for correct and incorrect trials. (A). Category information (PEV) for correct trials and incorrect trials for alpha/beta (left), gamma (middle), and spiking (right) under low-abstractness. Horizontal grey bars on the x-axis represented significant PEV differences between correct- and incorrect conditions (p<0.05, two-sample t-test). Error bar: mean ± SE, n = 51. (B). Cumulative difference in category information (correct minus incorrect) across time for each signal type. *** p<0.001, n.s.: not significant, one-sample t-test for each signal type and two-sample t-test for across signals. Error bar: mean ± SE, n = 51. (C-D). Same as (A-B) but under high-abstractness.

**Figure S8.**
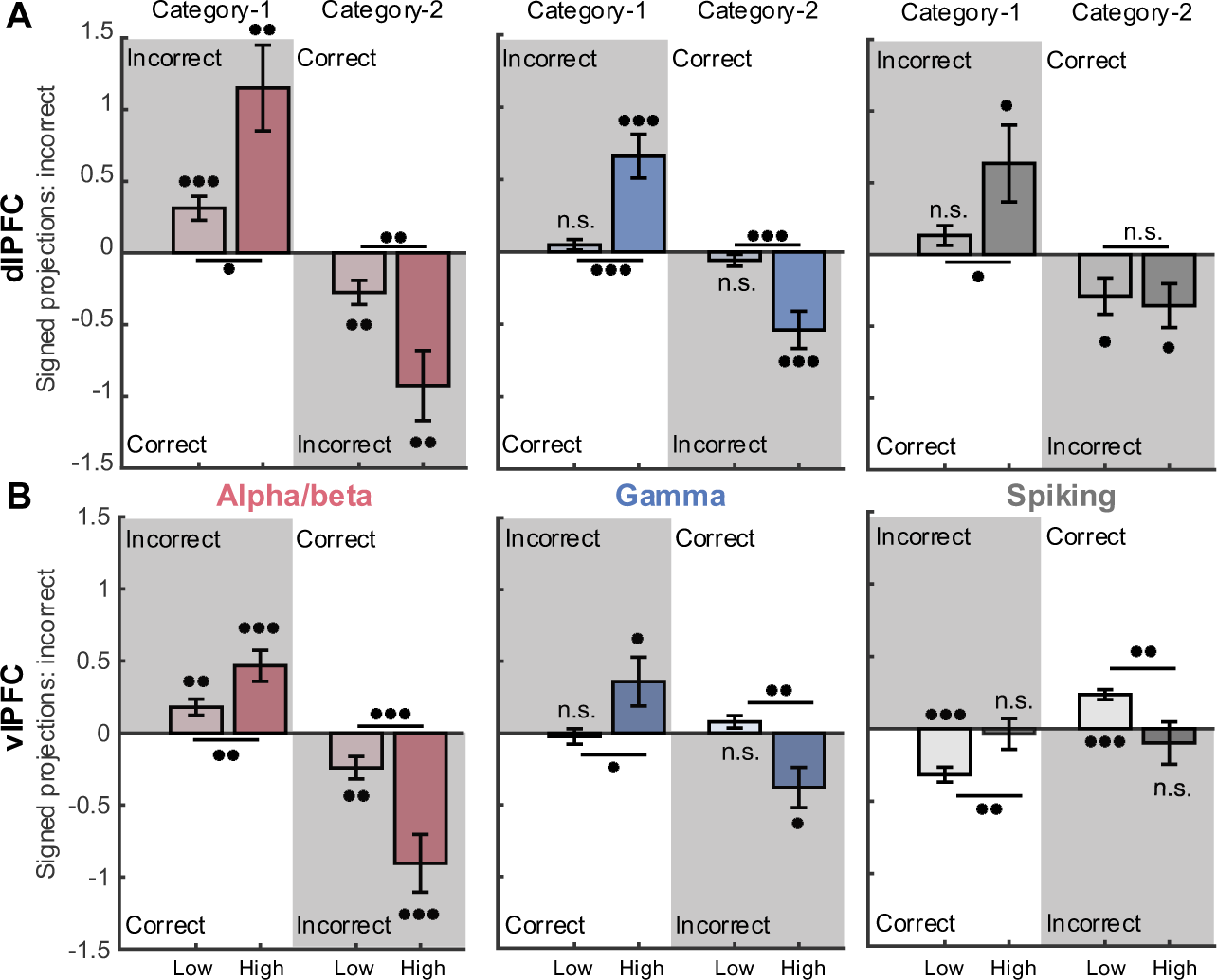
Signed projections of incorrect trials onto the decision axis for Category Task. (A). Signed projections of incorrect trials on the category axis of each category under low- and high- abstractness conditions, shown for alpha/beta (left), gamma (middle), and spiking (right) in dlPFC. *** p<0.001, n.s.: not significant, one-sample t-test for each signal type and Wilcoxon rank-sum test for across signals. Error bar: mean ± SE, n = 51. (B). Same as (A) but for vlPFC.

**Figure S9.**
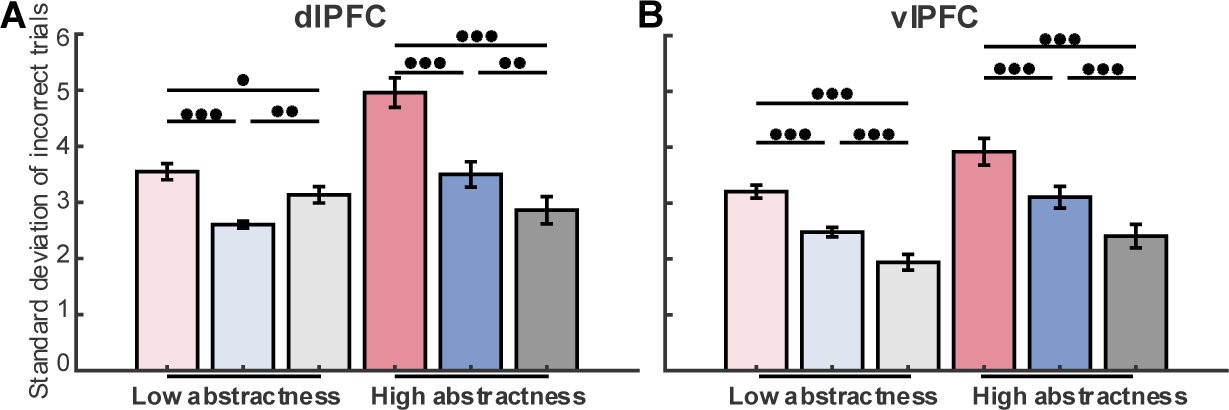
Standard deviation of the projections from incorrect trials for Category Task. (A). Standard deviation of the incorrect-trial projections for all signal types in dlPFC. *** p<0.001, **: p<0.01, *: p<0.05, one-sample t-test for each signal type and Wilcoxon rank-sum test for across signals. Error bar: mean ± SE, n = 51. (B). Same as (A) but for vlPFC. (C).

## Notes

### Competing Interest Statement

The authors have declared no competing interest.

### Summary of Updates

Corrected the missing word "information" in the Abstract.

## References

1. Lundqvist, M., Brincat, S.L., Rose, J., Warden, M.R., Buschman, T.J., Miller, E.K., and Herman, P. (2023). Working memory control dynamics follow principles of spatial computing. Nat Commun 14, 1429. 10.1038/s41467-023-36555-4.

2. Lundqvist, M., Miller, E.K., Nordmark, J., Liljefors, J., and Herman, P. (2024). Beta: bursts of cognition. Trends in Cognitive Sciences 28, 662–676. 10.1016/j.tics.2024.03.010.

3. Panichello, M.F., and Buschman, T.J. (2021). Shared mechanisms underlie the control of working memory and attention. Nature 592, 601–605. 10.1038/s41586-021-03390-w.

4. Sani, O.G., Abbaspourazad, H., Wong, Y.T., Pesaran, B., and Shanechi, M.M. (2021). Modeling behaviorally relevant neural dynamics enabled by preferential subspace identification. Nat Neurosci 24, 140–149. 10.1038/s41593-020-00733-0.

5. Duncan, J., and Miller, E.K. (2002). Cognitive focus through adaptive neural coding in the primate prefrontal cortex. In Principles of frontal lobe function (Oxford University Press), pp. 278–291. 10.1093/acprof:oso/9780195134971.003.0018.

6. Rigotti, M., Barak, O., Warden, M.R., Wang, X.-J., Daw, N.D., Miller, E.K., and Fusi, S. (2013). The importance of mixed selectivity in complex cognitive tasks. Nature 497, 585–590. 10.1038/nature12160.

7. Tye, K.M., Miller, E.K., Taschbach, F.H., Benna, M.K., Rigotti, M., and Fusi, S. (2024). Mixed selectivity: Cellular computations for complexity. Neuron 112, 2289–2303. 10.1016/j.neuron.2024.04.017.

8. Fusi, S., Miller, E.K., and Rigotti, M. (2016). Why neurons mix: high dimensionality for higher cognition. Current Opinion in Neurobiology 37, 66–74. 10.1016/j.conb.2016.01.010.

9. Warden, M.R., and Miller, E.K. (2010). Task-Dependent Changes in Short-Term Memory in the Prefrontal Cortex. J. Neurosci. 30, 15801–15810. 10.1523/JNEUROSCI.1569-10.2010.

10. Bhattacharya, S., Brincat, S.L., Lundqvist, M., and Miller, E.K. (2022). Traveling waves in the prefrontal cortex during working memory. PLoS Comput Biol 18, e1009827. 10.1371/journal.pcbi.1009827.

11. Kozachkov, L., Tauber, J., Lundqvist, M., Brincat, S.L., Slotine, J.-J., and Miller, E.K. (2022). Robust and brain-like working memory through short-term synaptic plasticity. PLOS Computational Biology 18, e1010776. 10.1371/journal.pcbi.1010776.

12. Loonis, R.F., Brincat, S.L., Antzoulatos, E.G., and Miller, E.K. (2017). A Meta-Analysis Suggests Different Neural Correlates for Implicit and Explicit Learning. Neuron 96, 521–534.e7. 10.1016/j.neuron.2017.09.032.

13. Wutz, A., Loonis, R., Roy, J.E., Donoghue, J.A., and Miller, E.K. (2018). Different Levels of Category Abstraction by Different Dynamics in Different Prefrontal Areas. Neuron 97, 716–726.e8. 10.1016/j.neuron.2018.01.009.

14. Warden, M.R., and Miller, E.K. (2007). The Representation of Multiple Objects in Prefrontal Neuronal Delay Activity. Cerebral Cortex 17, i41–i50. 10.1093/cercor/bhm070.

15. Donoghue, T., Haller, M., Peterson, E.J., Varma, P., Sebastian, P., Gao, R., Noto, T., Lara, A.H., Wallis, J.D., Knight, R.T., et al. (2020). Parameterizing neural power spectra into periodic and aperiodic components. Nat Neurosci 23, 1655–1665. 10.1038/s41593-020-00744-x.

16. Lundqvist, M., Herman, P., Warden, M.R., Brincat, S.L., and Miller, E.K. (2018). Gamma and beta bursts during working memory readout suggest roles in its volitional control. Nat Commun 9, 394. 10.1038/s41467-017-02791-8.

17. Petrides, M. (2005). Lateral prefrontal cortex: architectonic and functional organization. Philosophical Transactions of the Royal Society B: Biological Sciences 360, 781–795. 10.1098/rstb.2005.1631.

18. Mitchell, D.G.V., Rhodes, R.A., Pine, D.S., and Blair, R.J.R. (2008). The contribution of ventrolateral and dorsolateral prefrontal cortex to response reversal. Behavioural Brain Research 187, 80–87. 10.1016/j.bbr.2007.08.034.

19. Yamagata, T., Nakayama, Y., Tanji, J., and Hoshi, E. (2012). Distinct Information Representation and Processing for Goal-Directed Behavior in the Dorsolateral and Ventrolateral Prefrontal Cortex and the Dorsal Premotor Cortex. J. Neurosci. 32, 12934–12949. 10.1523/JNEUROSCI.2398-12.2012.

20. Moran, P.A.P. (1950). Notes on Continuous Stochastic Phenomena. Biometrika 37, 17–23. 10.2307/2332142.

21. Bullock, K.R., Pieper, F., Sachs, A.J., and Martinez-Trujillo, J.C. (2017). Visual and presaccadic activity in area 8Ar of the macaque monkey lateral prefrontal cortex. Journal of Neurophysiology 118, 15–28. 10.1152/jn.00278.2016.

22. Zhang, H., Watrous, A.J., Patel, A., and Jacobs, J. (2018). Theta and Alpha Oscillations Are Traveling Waves in the Human Neocortex. Neuron 98, 1269–1281.e4. 10.1016/j.neuron.2018.05.019.

23. Cohen, M.R., and Maunsell, J.H.R. (2010). A Neuronal Population Measure of Attention Predicts Behavioral Performance on Individual Trials. J. Neurosci. 30, 15241–15253. 10.1523/JNEUROSCI.2171-10.2010.

24. Buschman, T.J., Denovellis, E.L., Diogo, C., Bullock, D., and Miller, E.K. (2012). Synchronous Oscillatory Neural Ensembles for Rules in the Prefrontal Cortex. Neuron 76, 838–846. 10.1016/j.neuron.2012.09.029.

25. Richter, C.G., Coppola, R., and Bressler, S.L. (2018). Top-down beta oscillatory signaling conveys behavioral context in early visual cortex. Sci Rep 8, 6991. 10.1038/s41598-018-25267-1.

26. Salazar, R.F., Dotson, N.M., Bressler, S.L., and Gray, C.M. (2012). Content-Specific Fronto-Parietal Synchronization During Visual Working Memory. Science 338, 1097–1100. 10.1126/science.1224000.

27. Hoffman, S.J., Dotson, N.M., Lima, V., and Gray, C.M. (2024). The Primate Cortical LFP Exhibits Multiple Spectral and Temporal Gradients and Widespread Task-Dependence During Visual Short-Term Memory. Journal of Neurophysiology, jn.00264.2023. 10.1152/jn.00264.2023.

28. Lundqvist, M., Rose, J., Herman, P., Brincat, S.L., Buschman, T.J., and Miller, E.K. (2016). Gamma and Beta Bursts Underlie Working Memory. Neuron 90, 152–164. 10.1016/j.neuron.2016.02.028.

29. Ungerleider, L.G., Gaffan, D., and Pelak, V.S. (1989). Projections from inferior temporal cortex to prefrontal cortex via the uncinate fascicle in rhesus monkeys. Exp Brain Res 76, 473–484. 10.1007/BF00248903.

30. Sakagami, M., and Pan, X. (2007). Functional role of the ventrolateral prefrontal cortex in decision making. Current Opinion in Neurobiology 17, 228–233. 10.1016/j.conb.2007.02.008.

31. Hasan, A., Galea, J.M., Casula, E.P., Falkai, P., Bestmann, S., and Rothwell, J.C. (2013). Muscle and Timing-specific Functional Connectivity between the Dorsolateral Prefrontal Cortex and the Primary Motor Cortex. J Cogn Neurosci 25, 558–570. 10.1162/jocn_a_00338.

32. Morris, R.W., Dezfouli, A., Griffiths, K.R., and Balleine, B.W. (2014). Action-value comparisons in the dorsolateral prefrontal cortex control choice between goal-directed actions. Nat Commun 5, 4390. 10.1038/ncomms5390.

33. Bollimunta, A., Chen, Y., Schroeder, C.E., and Ding, M. (2008). Neuronal Mechanisms of Cortical Alpha Oscillations in Awake-Behaving Macaques. J. Neurosci. 28, 9976–9988. 10.1523/JNEUROSCI.2699-08.2008.

34. Siegel, M., Donner, T.H., and Engel, A.K. (2012). Spectral fingerprints of large-scale neuronal interactions. Nat Rev Neurosci 13, 121–134. 10.1038/nrn3137.

35. Haegens, S., Nácher, V., Luna, R., Romo, R., and Jensen, O. (2011). α-Oscillations in the monkey sensorimotor network influence discrimination performance by rhythmical inhibition of neuronal spiking. Proceedings of the National Academy of Sciences 108, 19377–19382. 10.1073/pnas.1117190108.

36. Jensen, O. (2024). Distractor inhibition by alpha oscillations is controlled by an indirect mechanism governed by goal-relevant information. Commun Psychol 2, 36. 10.1038/s44271-024-00081-w.

37. Klimesch, W. (2012). Alpha-band oscillations, attention, and controlled access to stored information. Trends in Cognitive Sciences 16, 606–617. 10.1016/j.tics.2012.10.007.

38. Foxe, J.J., and Snyder, A.C. (2011). The Role of Alpha-Band Brain Oscillations as a Sensory Suppression Mechanism during Selective Attention. Front. Psychol. 2. 10.3389/fpsyg.2011.00154.

39. Meyer, L., Obleser, J., and Friederici, A.D. (2013). Left parietal alpha enhancement during working memory-intensive sentence processing. Cortex 49, 711–721. 10.1016/j.cortex.2012.03.006.

40. Bonnefond, M., and Jensen, O. (2012). Alpha Oscillations Serve to Protect Working Memory Maintenance against Anticipated Distracters. Current Biology 22, 1969–1974. 10.1016/j.cub.2012.08.029.

41. Jensen, O., and Mazaheri, A. (2010). Shaping Functional Architecture by Oscillatory Alpha Activity: Gating by Inhibition. Front. Hum. Neurosci. 4. 10.3389/fnhum.2010.00186.

42. Miller, E.K., Lundqvist, M., and Bastos, A.M. (2018). Working Memory 2.0. Neuron 100, 463–475. 10.1016/j.neuron.2018.09.023.

43. Jokisch, D., and Jensen, O. (2007). Modulation of Gamma and Alpha Activity during a Working Memory Task Engaging the Dorsal or Ventral Stream. J. Neurosci. 27, 3244–3251. 10.1523/JNEUROSCI.5399-06.2007.

44. Haegens, S., Nácher, V., Hernández, A., Luna, R., Jensen, O., and Romo, R. (2011). Beta oscillations in the monkey sensorimotor network reflect somatosensory decision making. Proceedings of the National Academy of Sciences 108, 10708–10713. 10.1073/pnas.1107297108.

45. Ede, F. van, Lange, F. de, Jensen, O., and Maris, E. (2011). Orienting Attention to an Upcoming Tactile Event Involves a Spatially and Temporally Specific Modulation of Sensorimotor Alpha- and Beta-Band Oscillations. J. Neurosci. 31, 2016–2024. 10.1523/JNEUROSCI.5630-10.2011.

46. Spaak, E., Bonnefond, M., Maier, A., Leopold, D.A., and Jensen, O. (2012). Layer-Specific Entrainment of Gamma-Band Neural Activity by the Alpha Rhythm in Monkey Visual Cortex. Current Biology 22, 2313–2318. 10.1016/j.cub.2012.10.020.

47. van Kerkoerle, T., Self, M.W., Dagnino, B., Gariel-Mathis, M.-A., Poort, J., van der Togt, C., and Roelfsema, P.R. (2014). Alpha and gamma oscillations characterize feedback and feedforward processing in monkey visual cortex. Proceedings of the National Academy of Sciences 111, 14332– 14341. 10.1073/pnas.1402773111.

48. Bastos, A.M., Lundqvist, M., Waite, A.S., Kopell, N., and Miller, E.K. (2020). Layer and rhythm specificity for predictive routing. Proceedings of the National Academy of Sciences 117, 31459– 31469. 10.1073/pnas.2014868117.

49. Womelsdorf, T., Valiante, T.A., Sahin, N.T., Miller, K.J., and Tiesinga, P. (2014). Dynamic circuit motifs underlying rhythmic gain control, gating and integration. Nat Neurosci 17, 1031–1039. 10.1038/nn.3764.

50. Hamm, J.P., and Yuste, R. (2016). Somatostatin Interneurons Control a Key Component of Mismatch Negativity in Mouse Visual Cortex. Cell Reports 16, 597–604. 10.1016/j.celrep.2016.06.037.

51. Chen, G., Zhang, Y., Li, X., Zhao, X., Ye, Q., Lin, Y., Tao, H.W., Rasch, M.J., and Zhang, X. (2017). Distinct Inhibitory Circuits Orchestrate Cortical *beta* and *gamma* Band Oscillations. Neuron 96, 1403–1418.e6. 10.1016/j.neuron.2017.11.033.

52. Van Derveer, A.B., Bastos, G., Ferrell, A.D., Gallimore, C.G., Greene, M.L., Holmes, J.T., Kubricka, V., Ross, J.M., and Hamm, J.P. (2021). A Role for Somatostatin-Positive Interneurons in Neuro-Oscillatory and Information Processing Deficits in Schizophrenia. Schizophrenia Bulletin 47, 1385– 1398. 10.1093/schbul/sbaa184.

53. Lee, B., Shin, D., Gross, S.P., and Cho, K.-H. (2018). Combined Positive and Negative Feedback Allows Modulation of Neuronal Oscillation Frequency during Sensory Processing. Cell Reports 25, 1548–1560.e3. 10.1016/j.celrep.2018.10.029.

54. Domhof, J.W.M., and Tiesinga, P.H.E. (2021). Flexible Frequency Switching in Adult Mouse Visual Cortex Is Mediated by Competition Between Parvalbumin and Somatostatin Expressing Interneurons. Neural Computation 33, 926–966. 10.1162/neco_a_01369.

55. Zhao, S., Zhou, J., Zhang, Y., and Wang, D.-H. (2023). γ And β Band Oscillation in Working Memory Given Sequential or Concurrent Multiple Items: A Spiking Network Model. eNeuro 10. 10.1523/ENEURO.0373-22.2023.

56. Makino, H., and Komiyama, T. (2015). Learning enhances the relative impact of top-down processing in the visual cortex. Nat Neurosci 18, 1116–1122. 10.1038/nn.4061.

57. Studer, F., Zeppillo, T., and Barkat, T.R. (2024). Top-down inputs are controlled by somatostatin-expressing interneurons during associative learning. Preprint at bioRxiv, 10.1101/2024.12.18.629098 https://doi.org/10.1101/2024.12.18.629098.

58. Fries, P., Womelsdorf, T., Oostenveld, R., and Desimone, R. (2008). The Effects of Visual Stimulation and Selective Visual Attention on Rhythmic Neuronal Synchronization in Macaque Area V4. J. Neurosci. 28, 4823–4835. 10.1523/JNEUROSCI.4499-07.2008.

59. Ray, S., and Maunsell, J.H.R. (2011). Different Origins of Gamma Rhythm and High-Gamma Activity in Macaque Visual Cortex. PLOS Biology 9, e1000610. 10.1371/journal.pbio.1000610.

60. Michalareas, G., Vezoli, J., van Pelt, S., Schoffelen, J.-M., Kennedy, H., and Fries, P. (2016). Alpha-Beta and Gamma Rhythms Subserve Feedback and Feedforward Influences among Human Visual Cortical Areas. Neuron 89, 384–397. 10.1016/j.neuron.2015.12.018.

61. Womelsdorf, T., Fries, P., Mitra, P.P., and Desimone, R. (2006). Gamma-band synchronization in visual cortex predicts speed of change detection. Nature 439, 733–736. 10.1038/nature04258.

62. Siegel, M., Donner, T.H., Oostenveld, R., Fries, P., and Engel, A.K. (2008). Neuronal Synchronization along the Dorsal Visual Pathway Reflects the Focus of Spatial Attention. Neuron 60, 709–719. 10.1016/j.neuron.2008.09.010.

63. Cromer, J.A., Roy, J.E., and Miller, E.K. (2010). Representation of Multiple, Independent Categories in the Primate Prefrontal Cortex. Neuron 66, 796–807. 10.1016/j.neuron.2010.05.005.

64. Antzoulatos, E.G., and Miller, E.K. (2016). Synchronous beta rhythms of frontoparietal networks support only behaviorally relevant representations. eLife 5, e17822. 10.7554/eLife.17822.

65. Brincat, S.L., Donoghue, J.A., Mahnke, M.K., Kornblith, S., Lundqvist, M., and Miller, E.K. (2021). Interhemispheric transfer of working memories. Neuron 109, 1055–1066.e4. 10.1016/j.neuron.2021.01.016.

66. Antzoulatos, E.G., and Miller, E.K. (2011). Differences between Neural Activity in Prefrontal Cortex and Striatum during Learning of Novel Abstract Categories. Neuron 71, 243–249. 10.1016/j.neuron.2011.05.040.

67. Tallon-Baudry, C., and Bertrand, O. (1999). Oscillatory gamma activity in humans and its role in object representation. Trends in Cognitive Sciences 3, 151–162. 10.1016/S1364-6613(99)01299-1.

68. Siegel, M., Warden, M.R., and Miller, E.K. (2009). Phase-dependent neuronal coding of objects in short-term memory. Proc. Natl. Acad. Sci. U.S.A. 106, 21341–21346. 10.1073/pnas.0908193106.

69. Watson, B.O., Ding, M., and Buzsáki, G. (2018). Temporal coupling of field potentials and action potentials in the neocortex. Eur J of Neuroscience 48, 2482–2497. 10.1111/ejn.13807.

70. Olejnik, S., and Algina, J. (2003). Generalized Eta and Omega Squared Statistics: Measures of Effect Size for Some Common Research Designs. Psychological Methods 8, 434–447. 10.1037/1082-989X.8.4.434.

71. Chen, Z., and Padmanabhan, K. (2022). Top-down feedback enables flexible coding strategies in the olfactory cortex. Cell Reports 38, 110545. 10.1016/j.celrep.2022.110545.

72. Whiteway, M.R., Averbeck, B., and Butts, D.A. (2020). A latent variable approach to decoding neural population activity. Preprint at bioRxiv, 10.1101/2020.01.06.896423 https://doi.org/10.1101/2020.01.06.896423.

73. Calderini, M., and Thivierge, J.-P. (2021). Estimating Fisher discriminant error in a linear integrator model of neural population activity. The Journal of Mathematical Neuroscience 11, 6. 10.1186/s13408-021-00104-4.

74. Buschman, T.J., Siegel, M., Roy, J.E., and Miller, E.K. (2011). Neural substrates of cognitive capacity limitations. Proceedings of the National Academy of Sciences 108, 11252–11255. 10.1073/pnas.1104666108.

75. Gyurkovics, M., Clements, G.M., Low, K.A., Fabiani, M., and Gratton, G. (2022). Stimulus-Induced Changes in 1/ *f* -like Background Activity in EEG. J. Neurosci. 42, 7144–7151. 10.1523/JNEUROSCI.0414-22.2022.

